# A Rapid and Universal Pipeline for High-Resolution GPCR Structure Determination through *In Silico* Construct Optimization and de novo Protein Design

**DOI:** 10.64898/2026.04.02.716066

**Authors:** Asato Kojima, Kouki Kawakami, Naoya Kobayashi, Kazuhiro Kobayashi, Toshiki E. Matsui, Kohei Uemoto, Yuzhong Gu, Masahiro Fukuda, Hideaki E. Kato

## Abstract

G protein-coupled receptors (GPCRs) are critical regulators of human physiology and major drug targets. Although structural studies have provided valuable insights, determining GPCR structures remains challenging, especially for inactive state receptors. Recent advances in cryo-electron microscopy (cryo-EM) have enabled structural determination of small GPCRs by using fusion partner proteins and binders to increase molecular weight. However, current methods require extensive experimental screening of fusion constructs. Widely adopted strategies, such as BRIL-Fab complexes, also face limitations due to inherent flexibility. Here, we introduce a streamlined and universal pipeline that integrates an *in silico* fusion construct screening program, NOAH (NOAH: NOn-experimental, AI-assisted High-throughput construct screening), with a *de novo* designed fusion protein called ARK1 (ARtificially-designed fiducial marKer). We validate the efficacy of NOAH by determining the structures of the vasopressin V2 receptor (V2R) bound to the clinical antagonist tolvaptan and the partial agonist OPC51803, as well as the bradykinin B2 receptor (B2R) bound to the clinical antagonist icatibant, thereby elucidating their activation and deactivation mechanisms. Furthermore, we demonstrate the capability of NOAH-ARK1 by solving the tolvaptan-bound V2R structure at higher resolution and showcase the method’s versatility by determining the structure of lysophosphatidic acid receptor 2 (LPA2) bound to the antagonist Ki16425. This approach eliminates the need for time-consuming and labor-intensive construct optimization, providing a rapid and widely applicable solution for high-resolution GPCR structure determination and drug discovery.

## Introduction

G protein-coupled receptors (GPCRs) constitute the largest family of membrane proteins in the human genome, encompassing over 800 genes^1^. These seven-transmembrane receptors detect a wide range of extracellular signals, including light, odorants, hormones, neurotransmitters, and lipids, and modulate numerous physiological processes via their signaling partners, such as G proteins, G protein-coupled receptor kinases (GRKs), and arrestins. Consequently, GPCRs have emerged as pivotal drug targets, with nearly 35% of FDA-approved drugs acting on them^2^. To advance our molecular understanding and facilitate GPCR-targeted drug development, numerous structures have been solved in both ligand-bound and unbound states.

Historically, GPCR structures were elucidated by X-ray crystallography. Despite their inherent instability and small size, several innovations, particularly the lipidic cubic phase (LCP) crystallization method and fusion protein incorporation into intracellular loop 3 (ICL3) of receptors, have addressed these challenges^3^. Initially, T4 lysozyme was employed as a fusion partner; subsequently, other small soluble proteins, including thermostabilized apocytochrome b562 (BRIL), rubredoxin, flavodoxin, xylanase, and glycogen synthase (PGS), have also been utilized^3–7^. However, screening for crystallizable constructs and optimizing crystallization conditions remain a time-consuming and labor-intensive process, sometimes requiring years to solve a single structure^8^.

The advent of cryo-electron microscopy (cryo-EM) has significantly advanced the field by eliminating the need for crystals and large amounts of protein, markedly accelerating structure determination. Since the first cryo-EM structures of class B GPCR-Gs complexes in 2017^9,10^, over 200 unique GPCR-G protein complexes have been determined^11^. Nevertheless, successful image alignment in cryo-EM data processing requires protein targets to have both adequate molecular weight and well-defined structural features. Because class A GPCRs are relatively small (∼40 kDa) and often obscured by detergent micelles or lipid nanodiscs, early studies predominantly focused on agonist-bound, activated receptor-G protein complexes.

A comprehensive understanding of ligand-induced receptor activation and inactivation requires structural insights into both active and inactive GPCR states. Accordingly, several groups have pursued methods to efficiently determine antagonist-bound inactive GPCR structures using cryo-EM. In 2020, the first GPCR structural determination by cryo-EM without G proteins or other signaling partners was achieved by fusing a BRIL protein into ICL3 and employing both a Fab fragment and an anti-Fab nanobody^12^. Subsequent strategies included grafting ICL3 from the kappa-opioid receptor (KOR) with a nanobody targeting KOR-ICL3^13^, fusing PGS or calcineurin to the receptor^14,15^, and combining BRIL, an ALFA tag, anti-BRIL Fab, and a fusion protein of nanobodies targeting both BRIL-Fab and the ALFA tag^16^. However, these methods require extensive experimental screening to minimize flexibility between the receptor and its fusion partner, as well as separate expression and purification of multiple protein fragments, thereby significantly increasing both cost and time. Additionally, the widely used BRIL-Fab strategy often yields low-resolution structures, particularly in the receptor’s extracellular region (including the ligand-binding pocket), likely due to intrinsic flexibility of both BRIL and Fab components^4^.

To address these challenges, we developed an integrated pipeline leveraging recent advances in protein structure prediction and *de novo* protein design to efficiently determine GPCR structures without iterative experimental screening. First, we created NOAH (NOn-experimental, AI-assisted High-throughput construct screening), an *in silico* program generating optimal fusion constructs for cryo-EM by evaluating construct rigidity and stability. NOAH combines ColabFold predictions with structural assessments, including Dictionary of Secondary Structure in Proteins (DSSP) algorithm, predicted local-distance difference test (pLDDT) scores, “helix phase”, and hydrophobic-hydrophilic matching between fusion proteins and lipid bilayers. We demonstrated NOAH’s effectiveness by determining structures of vasopressin V2 receptor (V2R) bound to the clinical antagonist tolvaptan and bradykinin B2 receptor (B2R) bound to the clinical antagonist icatibant, elucidating their activation and deactivation mechanisms. Further validation was provided by determining the structure of V2R bound to the partial agonist OPC51803, confirming the method’s applicability to both antagonists and agonists. Next, we developed ARK1 (ARtificially-designed fiducial marKer), a *de novo*-designed fusion partner protein offering higher molecular weight and reduced flexibility compared to BRIL. The efficacy of the NOAH-ARK1 approach was further demonstrated by determining the structure of V2R bound to tolvaptan at higher resolution than its BRIL-fused counterpart, as well as the structure of lysophosphatidic acid receptor 2 (LPA2) bound to the antagonist Ki16425. We also evaluated the broader applicability of the NOAH-ARK1 pipeline to nearly 300 class A GPCRs, proposing a general strategy for semi-automated GPCR structure determination that eliminates labor-intensive experimental screening.

## Results

### Development of NOAH: An AI-Assisted Method for Selecting Optimal GPCR-Fiducial Chimeric Constructs

To reduce reliance on extensive experimental screening, we developed an *in silico* approach for selecting chimeric GPCR constructs, in which a GPCR is fused with a fiducial marker (fusion partner) protein. Previous studies have shown that a continuous helical connection between the target protein and the fiducial marker is critical for high-quality cryo-EM analysis using fusion strategies^14^. Consequently, our initial strategy was to use ColabFold^17^ to predict the structures of fiducial marker-fused GPCR constructs and then apply the DSSP algorithm^18^ to identify candidates featuring a continuous helical linkage (Fig. 1a).

**Fig. 1.**
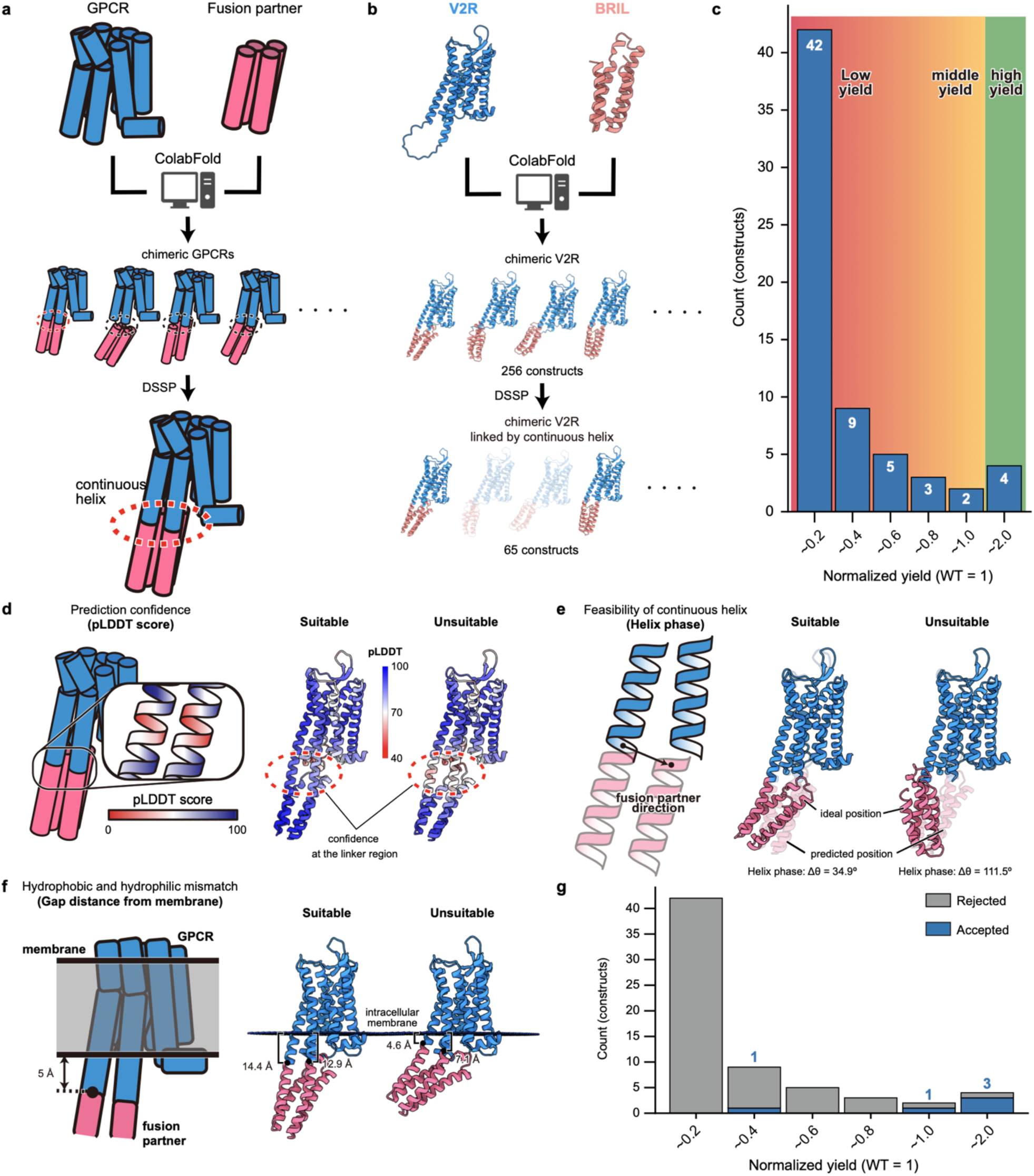
Development of NOAH, an AI-assisted pipeline for rational GPCR fusion construct selection. **a,** Initial workflow for *in silico* screening. ColabFold is employed to predict structural models of GPCR-fusion partner chimeras, followed by DSSP analysis, which selects constructs featuring a continuous α-helical linkage between receptor and fusion partner (dashed red circles). **b,** Application of this two-step screening procedure to V2R-BRIL fusions. From an initial set of 256 models generated by ColabFold, DSSP filtering yielded 65 candidate constructs. **c,** Histogram showing the relative expression levels (FSEC peak heights normalized to wild-type V2R = 1) for the 65 candidate constructs. **d–f,** Additional orthogonal criteria incorporated into NOAH to identify high-expression constructs. **d,** ColabFold confidence scores (pLDDT) along the TM5 and TM6 linker region; **e,** “Helix phase”, a geometric parameter evaluating the alignment between the C-terminal end of the fusion partner and the N-terminal end of TM6 of the receptor, essential for a straight α-helical connection between the receptor and fusion partner; **f,** Minimum distance between the fusion partner and the membrane boundary, required to exceed 5 Å to prevent hydrophophilic/hydrophobic mismatch. **g,** Distribution of relative expression levels after application of all four selection criteria (DSSP, pLDDT scores, helix phase, and membrane boundary gap).

To validate this method, we selected the vasopressin V2 receptor (V2R), a prototypical GPCR, as a model system. Initially, we designed 256 candidate V2R-BRIL constructs by fusing the N- and C-termini of BRIL to V2R at various positions within the TM5-ICL3-TM6 region (Extended Data Fig. 1a). Using ColabFold to predict the structures and DSSP to filter candidates, we reduced the candidate pool to 65 constructs with continuous helical linkers (Fig. 1b and Extended Data Fig. 1b).

To evaluate expression levels, we fused EGFP to the C-termini of these constructs and transiently expressed them in HEK293 cells. Fluorescence-detection size-exclusion chromatography (FSEC) analysis^19^ revealed that only approximately 6% of the constructs achieved expression levels comparable to or exceeding wild-type (WT) V2R, whereas 65% produced less than 20% of WT yields (Fig. 1c and Extended Data Fig. 1c). These results indicated that selection based solely on ColabFold and DSSP predictions is insufficient, necessitating additional yield-related criteria.

We hypothesized that the low expression yields observed might result from two primary factors. First, the enforced helical linkage between V2R and BRIL using suboptimal linkers may induce structural distortions and compromise stability, even when the predicted models exhibit continuous helices. Second, a hydrophobic-hydrophilic mismatch between BRIL and the lipid bilayer—an aspect not addressed in ColabFold predictions—could further widen the gap between the predicted and actual structures.

To mitigate these issues, we introduced three additional selection criteria. First, we assessed the accuracy of the predicted linker-region structure using the pLDDT score, which assigns lower scores to regions with less reliable secondary structures^20^ (Fig. 1d). Second, we imposed an angular constraint to ensure an optimal junction between the receptor and BRIL (Fig. 1e). Specifically, we treated the receptor and BRIL as rigid bodies by connecting the receptor’s TM5 C-terminus to the BRIL N-terminus and then evaluated the gap between the BRIL C-terminus and the receptor’s TM6 N-terminus. We defined this gap, evaluated as a function of the angle, as the “helix phase” (Fig. 1e and Extended Data Fig. 1d). If the BRIL C-terminal end is positioned too far from the receptor and if substantial deviation exists between this configuration and the ColabFold-predicted model, the structure may develop significant distortions that ultimately compromise the construct’s stability in practice, even if the predicted model exhibits continuous helices connecting TM5 to the first helix of BRIL and the last helix of BRIL to TM6. Third, to mitigate the hydrophobic-hydrophilic mismatch between BRIL and the lipid bilayer, we required the predicted membrane by PPM3^21^ and the fusion partner protein to maintain a minimum non-contact distance greater than 5 Å^22^ (Fig. 1f and Extended Data Fig. 1g).

Applying these three additional criteria to the 65 constructs preselected by ColabFold and DSSP further reduced the candidate pool: 29 constructs satisfied the first criterion (Extended Data Fig. 1c,e), 17 met the second (Extended Data Fig. 1c,f), and 27 fulfilled the third (Extended Data Fig. 1c,g), with only five constructs meeting all three criteria (Extended Data Fig. 1h-j). Notably, four of these five constructs exhibited expression levels comparable to or exceeding those of wild-type V2R, demonstrating that the selection process successfully enriched for constructs suitable for structural studies (Fig. 1g and Extended Data Fig. 1c,h-j).

We designated this integrated construct-selection method, which combines ColabFold and DSSP with the three additional criteria, as NOAH (NOn-experimental, AI-assisted High-throughput construct screening for structural analysis). Using V2R as a model system, NOAH efficiently reduced several hundred candidate constructs to a highly refined set of fewer than ten constructs suitable for experimental verification. Importantly, since these criteria are orthogonal, they likely represent the minimal necessary standards for construct selection in protein fusion strategies (Extended Data Fig. 1i, j).

### Determination of the antagonist-bound V2R-BRIL structure

Next, we evaluated whether the V2R-BRIL constructs selected using NOAH were suitable for single-particle cryo-EM analysis. Among the four constructs meeting our criteria, we selected the one with exhibiting a higher yield than WT V2R and minimal sequence truncation, and expressed it in HEK293 cells (Extended Data Fig. 2a). For structure determination of the antagonist-bound receptor, we solubilized and purified the receptor in the presence of the clinically used antagonist tolvaptan, complexed it with the anti-BRIL antibody SRP2070^23^, and vitrified the resulting assembly on cryo-EM grids (Extended Data Fig. 2b). Using a Titan Krios cryo-electron microscope, we imaged the vitrified sample and determined its structure at a nominal resolution of 3.0 Å, with well-defined density observed for most of the V2R-BRIL complex and for tolvaptan (Fig. 2a and Extended Data Fig. 2c-g). These results demonstrate the suitability of the NOAH-selected construct for cryo-EM analysis of antagonist-bound GPCRs. Furthermore, the interface between V2R and BRIL exhibited a continuous helix closely matching the ColabFold-predicted model, further validating our selection strategy (Fig. 2a and Extended Data Fig. 2h).

**Fig. 2.**
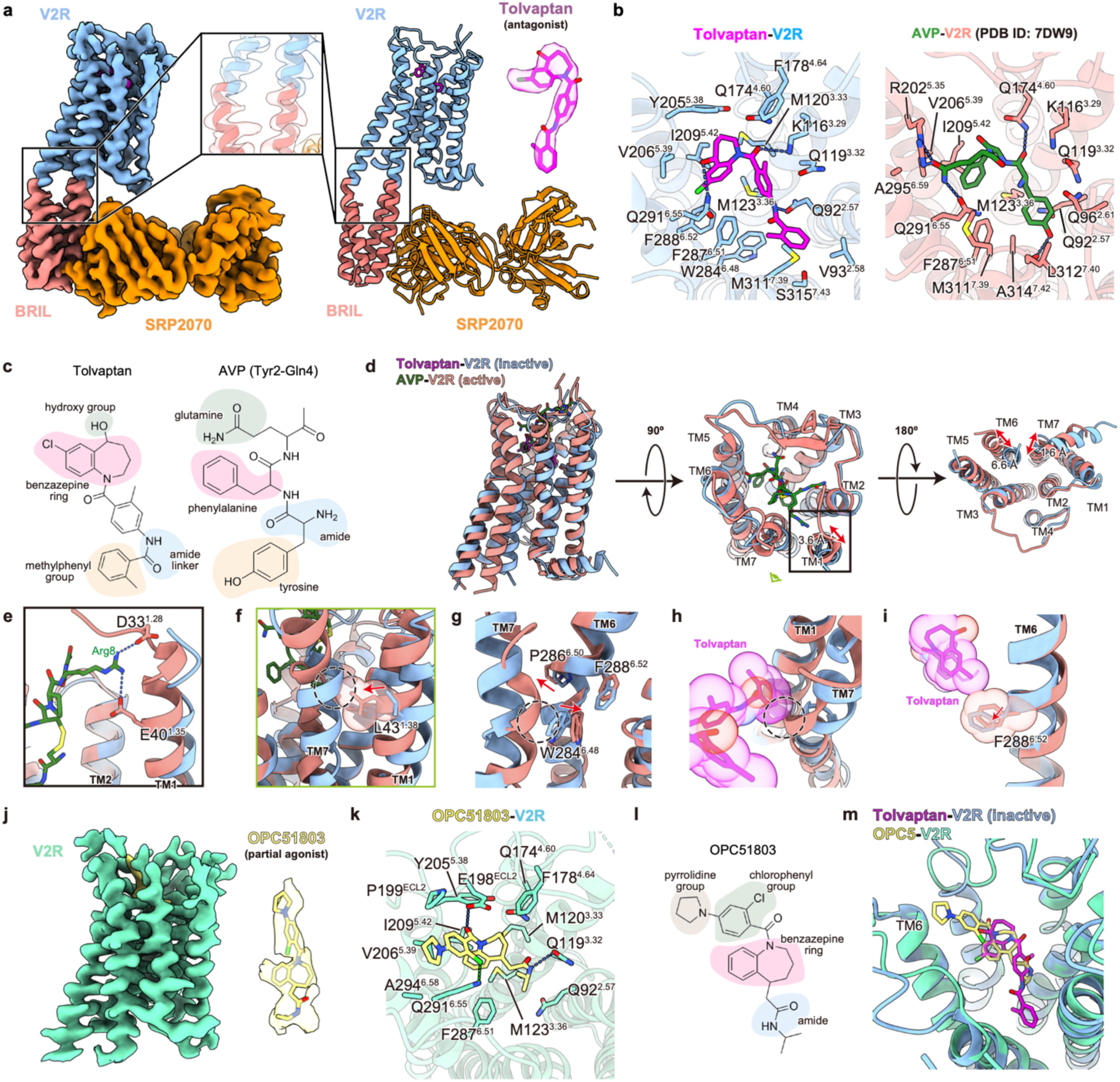
Structural analysis of V2R-BRIL bound to the clinical antagonist tolvaptan and partial agonist OPC5. **a,** Cryo-EM density map (left) and corresponding atomic model (right) of the tolvaptan-bound V2R-BRIL in complex with the anti-BRIL Fab SRP2070. Insets depict the continuous α-helical junction between V2R and BRIL (center) and the well-resolved density for tolvaptan (upper right). **b,** Ligand-binding pockets illustrating interactions of tolvaptan (this study, left) and the endogenous agonist arginine-vasopressin (AVP; PDB: 7DW9, right) with V2R. **c,** Chemical structures of tolvaptan and the Tyr2-Phe3-Gln4 tripeptide segment of AVP, with pharmacophoric elements: hydroxy/Gln4, benzazepine/Phe3, amide linker/amide, and methyl-phenyl/Tyr2. **d,** Comparison of receptor conformations in the tolvaptan-bound and the AVP-bound states, viewed parallel to the membrane (left), from the extracellular side (center; tolvaptan omitted), and from the intracellular side (right). **e,** Polar interaction network formed by Arg8 of AVP in the active-state structure. **f,** Partial unwinding of TM7 in the AVP-bound receptor caused by a steric clash with L43^1.38^ (red arrow). **g,** Outward movement of TM6 induced by the shift of partially unwound TM7. Movements of W284^6.48^ and P286^6.50^ are indicated by red arrows, and the steric clash between TM6 and TM7 is highlighted by a dashed black circle. **h,** Model illustrating how tolvaptan would sterically clash with TM7 in its active conformation. **i,** Conformational change of F288^6.52^ accommodating the benzazepine core of tolvaptan. **j,** Cryo-EM density of the partial agonist OPC5 bound to V2R (left), and overlay of the OPC5 density with the atomic model (right). **k,** Detailed interactions between OPC5 and residues lining the orthosteric pocket, highlighting hydrogen bonds (dashed lines) and hydrophobic contacts. **l,** Chemical structure of OPC5 with functional groups: pyrrolidine, chlorophenyl, benzazepine ring, and amide. **m,** Structural superposition of OPC5 and tolvaptan.

The structure reveals that tolvaptan interacts extensively with TM2-TM7 of V2R. Specifically, the hydroxy and carbonyl groups adjacent to its benzazepine ring form hydrogen bonds with K116^3.29^ and Q291^6.55^ (superscripts denote Ballesteros-Weinstein numbering^24^, while the amide linker connecting its two methylphenyl groups forms a hydrogen bond with Q92^2.57^ (Fig. 2b,c, left). Notably, these contacts partially mimic the Tyr2-Phe3-Gln4 motif binding of the endogenous V2R agonist arginine vasopressin (AVP) (Fig. 2b,c, right). In addition to hydrogen bonding, the benzazepine ring and methylphenyl groups of tolvaptan are stabilized by extensive hydrophobic or van der Waals interactions with residues in TM3 (Q119^3.32^, M120^3.33^, and M123^3.36^), TM5 (Y205^5.38^, V206^5.39^, and I209^5.42^), and TM6 (W284^6.48^, F287^6.51^, and F288^6.52^) as well as with residues in TM2 (Q92^2.57^ and V93^2.58^) and TM7 (M311^7.39^ and S315^7.43^) (Fig. 2b,c, left). These interactions, coupled with the aforementioned hydrogen bonds, underlie tolvaptan’s high potency (Ki = 0.43 ± 0.06 nM)^25^.

### Activation and deactivation mechanisms of V2R

To elucidate how tolvaptan stabilizes V2R in an inactive conformation, we compared our tolvaptan-bound V2R structure onto the previously reported AVP-bound active V2R structure^26^. TM2-TM5 align closely, whereas the extracellular ends of TM1 in the tolvaptan-bound receptor shift outward by 3.6 Å. Concomitantly, the intracellular end of TM6 moves inward by 6.6 Å and TM7 shifts outward by 1.6 Å, both characteristic hallmarks of an inactive GPCR (Fig. 2d).

To dissect the activation process, we compared the AVP-activated and tolvaptan-inactivated conformations. In the active state, AVP’s N-terminal eight residues penetrate the extracellular pocket, interacting with all TM helices (Extended Data Fig. 3a). Arg8 of AVP forms salt bridges with D33^1.28^ and E40^1.35^, drawing TM1 inward by 3.3 Å (Fig. 2e). This movement pushes L43^1.38^ into TM7, inducing partial unwinding of the helix (Fig. 2f). Additionally, Tyr2 of AVP forms a hydrogen bond with the backbone of L312^7.40^, stabilizing the new TM7 configuration (Extended Data Fig. 3b). The repositioned TM7 subsequently clashes with the toggle-switch residue W284^6.48^ (Fig. 2g), triggering a rotation and kink around P286^6.50^ that swings the intracellular end of TM6 outward, a signature of receptor activation.

In contrast, when tolvaptan is docked into the active state receptor, its methylphenyl substituents sterically clash with TM7, and its benzazepine core collides with F288^6.52^ in this conformation (Fig. 2h,i). These steric hindrances prevent both the inward shift of TM7 and the outward rotation of TM6, thereby locking the receptor in its inactive state and antagonizing activation.

### Determination of the agonist-bound V2R-BRIL structure

We next evaluated whether the NOAH-selected construct was suitable for solving the agonist-bound structure. Using OPC51803 (OPC5), a small-molecule partial agonist under clinical investigation, we performed cryo-EM analysis and obtained a high-resolution 3.3 Å map of the OPC5-bound V2R structure, with clear density for both V2R and OPC5 (Fig. 2j, Extended Data Fig. 4), confirming the construct’s suitability for structural analysis of agonist-bound receptors.

The OPC5-bound structure reveals fewer contacts with TM2-TM6 compared to AVP or tolvaptan (Fig. 2k,l). Remarkably, although OPC5 shares a backbone scaffold with tolvaptan, it binds the receptor in a ∼180° rotated orientation (Fig. 2m). In this orientation, the chlorophenyl group forms a halogen bond with Q291^6.55^, while its amide and carbonyl moieties engage Q119^3.32^ and E198^ECL2^ through hydrogen bonds. Additionally, the pyrrolidine, 2-chlorophenyl, and benzazepine rings establish hydrophobic or van der Waals interactions with Q92^2.57^, M120^3.33^, M123^3.36^, Q174^4.60^, F178^4.64^, P199^ECL2^, Y205^5.38^, V206^5.39^, I209^5.42^, F287^6.51^, and A294^6.58^ (Fig. 2k,l). These distinct ligand-receptor contacts likely underlie the pharmacological differences between OPC5 and tolvaptan. Notably, the fused BRIL protein locks V2R in an inactive-like conformation (Extended Data Fig. 3d); however, OPC5’s contacts with TM6 induce a subtle rotation and displacement of TM6 (Extended Data Fig. 3e). This suggests that OPC5 may activate V2R by directly promoting TM6 movement rather than rearranging TM1 and TM7, as observed in AVP-induced activation (Extended Data Fig. 3f).

### Determination of the antagonist-bound B2R-BRIL structure

To test NOAH’s applicability across class A GPCRs, we next chose the bradykinin B2 receptor (B2R), whose inactive conformation has not yet been determined. We generated 100 candidate constructs by inserting BRIL into the intracellular TM5-ICL3-TM6 segment (Extended Data Fig. 5a). ColabFold models of these constructs were first filtered by DSSP, reducing the pool to 15. We then removed any construct that failed to satisfy all three criteria— (1) a minimum pLDDT score, (2) helix-phase compatibility, and (3) sufficient separation between BRIL and the predicted membrane bilayer—leaving four viable constructs (Fig. 3a, Extended Data Fig. 5b-d). From these, we selected the variant whose sequence most closely matched that of WT B2R for experimental study (Extended Data Fig. 5e).

**Fig. 3.**
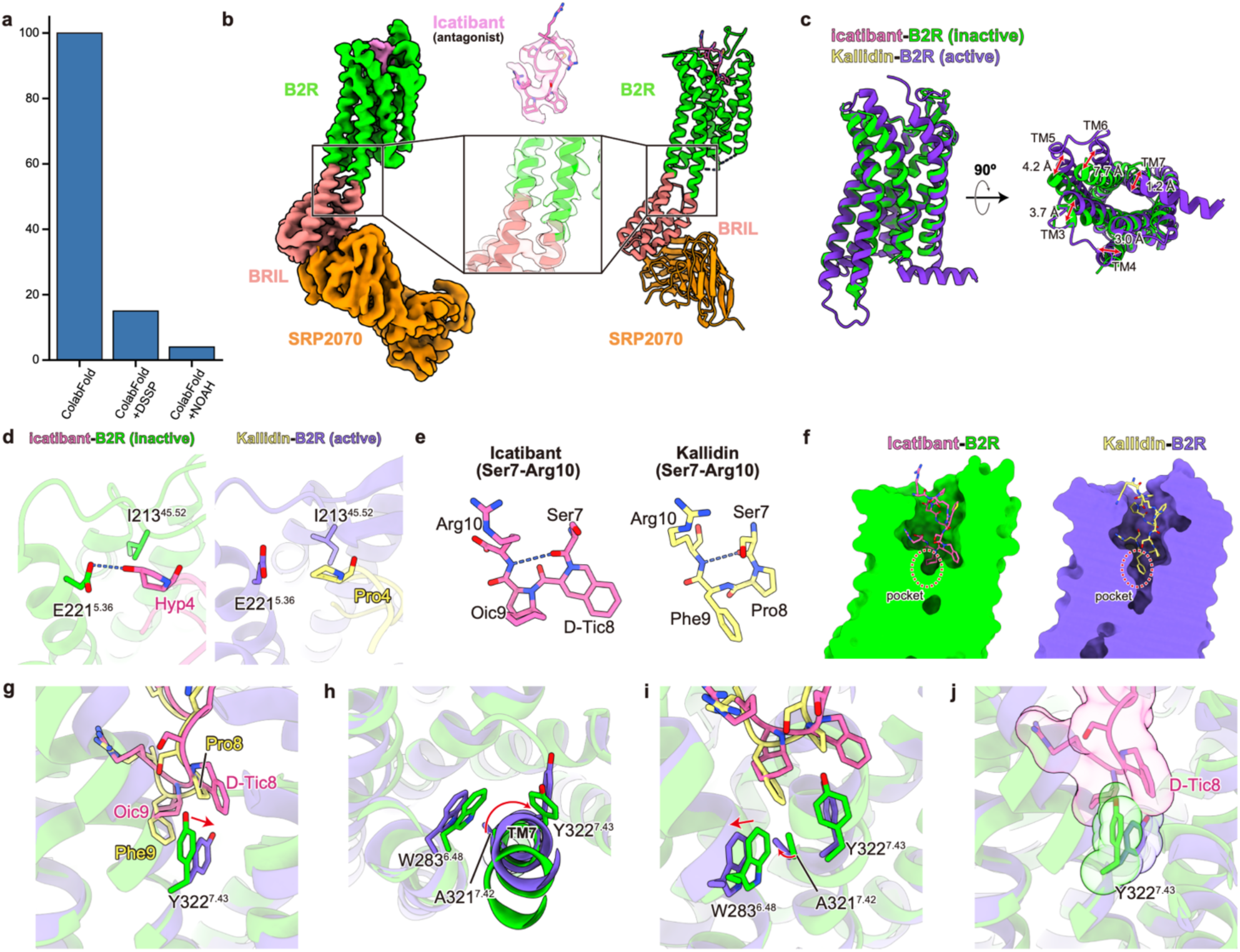
Structural analysis of B2R-BRIL bound to the clinical antagonist icatibant. **a,** Histogram illustrating the number of B2R-BRIL fusion constructs retained after successive *in silico* filtering steps: ColabFold alone, ColabFold + DSSP, and the complete NOAH pipeline. **b,** Cryo-EM density map (left) and corresponding atomic model (right) of icatibant-bound B2R-BRIL in complex with anti-BRIL Fab SRP2070. Insets highlight the continuous α-helical B2R-BRIL junction (center) and the density corresponding to icatibant (top). **c,** Structural superposition of inactive icatibant-bound B2R and active kallidin-bound B2R (PDB: 7F61), viewed parallel to the membrane (left) and from the intracellular side (right). **d,** Detailed interactions of Hyp4 in icatibant or Pro4 in kallidin with residues in B2R. **e,** Backbone traces of residues Ser7-Arg10 in icatibant and kallidin. Hydrogen bonds are indicated by dashed lines. **f,** Cross-sectional views of the icatibant- and kallidin-binding pockets in B2R. **g,** Structural comparison of Y322^7.43^ between kallidin- and icatibant-bound structures. Steric clash between Phe9 of kallidin and Y322^7.43^ forces Y322^7.43^ downward in the kallidin-bound structure. **h,** Clockwise rotation of TM7 (red arrow) induced by the movement of Y322^7.43^. **i,** Displacement of W283^6.48^ triggered by TM7 rotation via the movement of A321^7.42^. **j,** The bulky D-Tic8 residue of icatibant sterically blocks Y322^7.43^, preventing TM7 rotation and locking B2R in an inactive conformation.

This construct was expressed in HEK293 cells, solubilized and purified in the presence of the clinical antagonist icatibant, and complexed with the anti-BRIL antibody SRP2070 (Extended Data Fig. 5f). Single-particle cryo-EM on a Titan Krios microscope yielded a 3.7 Å reconstruction of the icatibant-bound B2R-BRIL complex, with clear density for both the transmembrane helices and the ligand (Fig. 3b, Extended Data Fig. 5g-k), enabling unambiguous model building. Relative to active-state structures bound to endogenous agonists bradykinin (RPPGFSPFR) or kallidin (KRPPGFSPFR)^27^, the icatibant-bound receptor adopts the canonical inactive conformation, characterized by an inward shift of TM6, closely matching our *in silico* predictions (Fig. 3c, Extended Data Fig. 5l).

Icatibant is different from the endogenous peptide kallidin at five positions—D-Arg1, Hyp4, Thi6, D-Tic8, and Oic9 (Extended Data Fig. 6a)—modifications designed to resist proteolysis and extend its pharmacological action^28^. To evaluate the roles of these substitutions, we compared the structures of B2R bound to icatibant and kallidin in detail. First, the N-terminal D-Arg1 of icatibant exhibits almost no interaction with adjacent residues, as indicated by weak density in the cryo-EM map (Extended Data Fig. 6b). This observation is consistent with the nearly identical affinities of bradykinin and kallidin for B2R^29^, suggesting that the substitution of D-Arg1 primarily serves to prevent proteolytic degradation. Additionally, the conformations of Arg2 and Thi6 in icatibant closely resemble those of Arg2 and Phe6 in kallidin, further indicating that these substitutions mainly confer resistance to degradation rather than enhancing receptor affinity (Extended Data Fig. 6c, d). Next, we focused on Hyp4 and found that its hydroxyl moiety forms additional hydrogen bonds with E221^5.36^ and R297^6.62^ (Fig. 3d), interactions absent with Pro4 in kallidin. This additional interaction rationalizes the higher affinity of icatibant, consistent with previous studies^30^. Finally, we analyzed the significance of the D-Tic8-Oic9 substitution. Comparing receptor-bound structures of kallidin and icatibant, we found that both peptides form a β-turn structure between residues 7-10, with remarkably similar conformations (Fig. 3e). However, due to the difference in size, Phe9 in kallidin penetrates deeper into the ligand-binding pocket compared to Oic9 in icatibant, interacting with and pushing Y322^7.43^ (Fig. 3f,g). This movement of Y322^7.43^ induces a clockwise rotation of TM7, with A321^7.42^ moving accordingly (Fig. 3h). A321^7.42^ then creates a steric clash with W283^6.48^, inducing the outward movement of TM6, which is required for G-protein coupling (Fig. 3i). In contrast, Oic9 in icatibant, due to its smaller size, cannot effectively push Y322^7.43^, and the bulky D-Tic8 sterically blocks the rearrangement of Y322^7.43^, preventing the TM7-TM6 activation cascade (Fig. 3j). Supporting this model, substitution of Pro8 in kallidin with a larger residue shifts the ligand’s pharmacological property from agonist to antagonist^31^. Thus, the movement of TM7 and TM6 triggered by Phe9 in kallidin plays a critical role in B2R activation, whereas the D-Tic8-Oic9 substitution in icatibant prevents activation by cooperatively blocking the movement of Y322^7.43^ (Extended Data Fig. 6e).

### Development of a *de novo* designed fusion partner, ARK1

NOAH construct screening significantly facilitates cryo-EM structural determination of GPCRs in both antagonist- and agonist-bound states. However, we recognized that the BRIL-Fab system has several limitations and identified opportunities for improvement. First, local resolution is typically highest in the BRIL-Fab region, while resolution in the GPCR region, particularly around the extracellular ligand-binding site, often remains lower, likely due to flexibility within BRIL, the Fab (Extended Data Fig. 7a), and at their interface (Extended Data Figs. 2e, 5i). Indeed, in some studies, extracellular binding sites have local resolutions of 4-5 Å despite overall map resolutions of 3.5 Å or better^32–34^. Second, antibody preparation (e.g., anti-BRIL Fab or NbFab) incurs substantial time and labor costs. Third, Fab binding can introduce orientation bias, degrading map quality (Extended Data Fig. 7b). For example, the V2R-BRIL complex without Fab exhibits less orientation bias than the V2R-BRIL-Fab assembly (although its smaller size prevented high-resolution map reconstruction) (Extended Data Fig. 7c-h).

To overcome these challenges, we sought a single, large, rigid fusion partner compatible with TM5/6 fusions. Since no natural proteins meet these requirements^4^, we turned to *de novo* design. *De novo* proteins are generally rigid and stable, often increasing the expression and stability of fusion constructs^35^. Thus, they appeared suitable for addressing the limitations of the BRIL-Fab system.

Using V2R as a model, we designed a ∼40 kDa protein—large enough to function as a robust fiducial marker for cryo-EM, yet tractable for *de novo* design. Although the BRIL-Fab ensemble has a combined mass of ∼60 kDa, Fab hinge flexibility reduces its effective rigid mass to roughly 35 kDa, suggesting that a rigid ∼40 kDa partner would meet these requirements.

The design principle is as follows (Fig. 4a). First, we generated two intracellular helices extending from TM5 and TM6 of V2R, each approximately 20 residues long, since excessively long helices can introduce orientation bias^36^. Next, these two helices served as nucleation points for two asymmetrical helical bundles consisting of six and seven helices (including the initial helices). This asymmetry provides distinct structural features, facilitating orientation assignment during cryo-EM analysis. Finally, we created a central bundle by inserting a helix between the left and right bundles to interlock the two peripheral bundles.

**Fig. 4.**
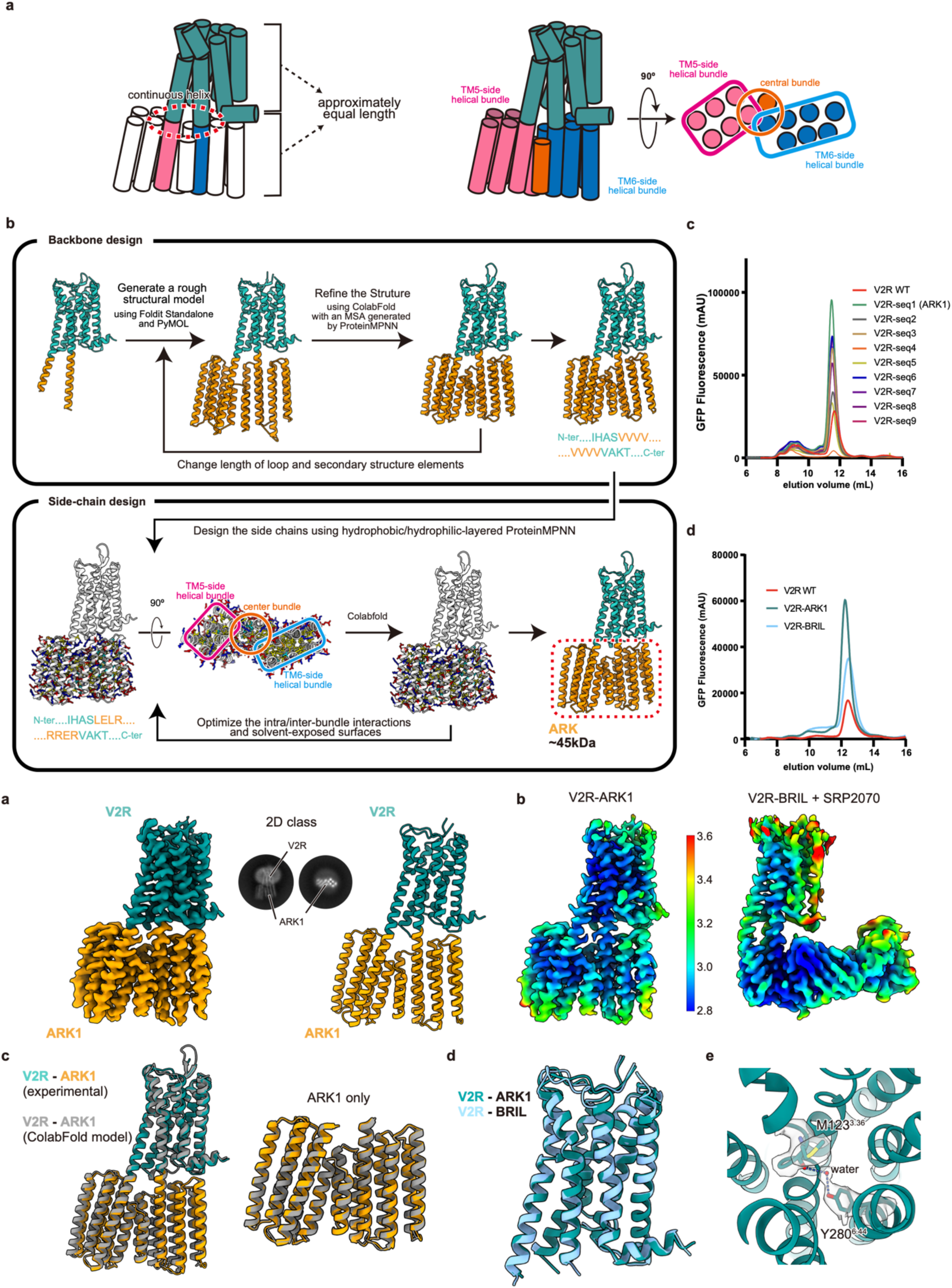
De novo design of ARK1, an ideal fiducial marker for cryo-EM analysis. **a,** Design principles for a rigid fiducial marker. ARK1 connects to the receptor through a single, continuous α-helix that exits TM5 and re-enters TM6 with approximately equal-lengths (left). The soluble portion folds into a compact, asymmetric three-helix bundle, providing high-contrast features for particle alignment (right). **b,** Two-stage computational pipeline for ARK1 design. Backbone design (top) began with a rough model created in Foldit/PyMOL, which was iteratively rebuilt and refined using ColabFold predictions guided by multiple-sequence alignments generated with ProteinMPNN. Sequence design (bottom) utilized a hydrophobic/hydrophilic-layered version of ProteinMPNN to optimize core packing and surface polarity, followed by targeted mutations to resolve residual mismatches while preserving high pLDDT scores. **c,** FSEC analysis of nine V2R-ARK fusion constructs (seq1-seq9). **d,** FSEC comparison of V2R-ARK1, wild-type V2R, and the V2R-BRIL fusion. **e,** Cryo-EM density map (left), corresponding atomic model (right), and representative 2D class average (center) of the tolvaptan-bound V2R-ARK1 complex. **f,** Local-resolution comparison between the V2R-ARK1 structure (left) and the V2R–BRIL structure in complex with SRP2070 Fab (right), as presented in Fig. 2. The local resolution around the receptor’s orthosteric pocket is significantly improved in the V2R-ARK1 structure. **g,** Structural superposition of the experimental V2R-ARK1 structure with the ColabFold prediction. **h,** Structural superposition of overall receptor backbones of V2R-ARK1 and V2R-BRIL. **i,** A water-mediated interaction between tolvaptan and V2R.

Following this principle, we initially built the core backbone in Foldit Standalone^37^ from a poly-Val peptide. We then employed our original protocols, named “BDP,” to refine the backbone (Kobayashi *et al.,* in preparation). Specifically, we generated a multiple sequence alignment (MSA) using ProteinMPNN^38^, predicted structures using ColabFold using the MSA as a guide, manually adjusted helices and loops in PyMOL, and iteratively refined the models (Fig. 4b, top). After refining the backbone structure, we generated 128 candidate sequences using Protein MPNN biased GCN-design, created predicted structures with ColabFold, and narrowed these down to nine candidates based on their pLDDT scores. We then optimized the hydrophobic core packing, inter-bundle interactions, and solvent-exposed surfaces to minimize orientation bias on cryo-EM grids (Fig. 4b, bottom). Each of these nine designed proteins was genetically fused to V2R, expressed in HEK293 cells, and evaluated using FSEC for peak intensity and monodispersity. Remarkably, all nine constructs showed monodisperse peaks, and eight constructs exhibited enhanced expression relative to WT V2R (Fig. 4c). Thus, we selected the candidate with the highest peak intensity, naming it ARK1 (ARtificially-designed fiducial marKer 1). Notably, the expression level of V2R-ARK1 was approximately four-fold and two-fold higher than those of WT V2R and V2R-BRIL, respectively (Fig. 4d).

### Determination of the antagonist-bound V2R-ARK1 structure

We next evaluated the suitability of the V2R-ARK1 construct for cryo-EM analysis. Samples were prepared using the same protocol applied to the tolvaptan-bound V2R-BRIL complex (Extended Data Fig. 2), and micrographs were acquired on a Titan Krios microscope (Extended Data Fig. 8). During image processing, the V2R-ARK1 construct displayed significantly reduced orientation bias, yielding an isotropic distribution of particle views (Extended Data Fig. 8g). From these data, we reconstructed a 2.98 Å resolution map of the tolvaptan-V2R-ARK1 assembly (Fig. 4e).

Notably, although the nominal resolution matched that of the V2R-BRIL structure, the V2R-ARK1 structure exhibited substantially improved local resolution around the receptor, particularly at the ligand-binding site, clearly revealing water molecules that facilitate receptor–ligand interactions. (Fig. 4f,g). The resulting structure closely aligned with our *in silico* predictions, displaying continuous helices connecting the receptor to ARK1 (Fig. 4h). A comparison of the receptor conformations in the V2R-ARK1 and V2R-BRIL structures revealed that their receptor folds are virtually indistinguishable, indicating that fusion with ARK1 (or BRIL) does not perturb the native V2R fold (Fig. 4i). Collectively, these results establish ARK1 as a novel fusion partner that enhances expression and improves map resolution, particularly at the orthosteric binding site and ligand, relative to the BRIL-Fab system.

### ARK-fusion construct design for general class A GPCRs

Having established ARK1’s effectiveness with V2R, we next explored whether the NOAH-ARK1 combination could be extended to generate ARK1-fusion constructs for cryo-EM analysis across the non-olfactory class A GPCR family. We applied four selection criteria—DSSP secondary-structure compatibility, pLDDT confidence, helix-phase alignment, and a minimum distance from the fusion partner to the lipid bilayer—to inactive-state predictions of 287 receptors (Fig. 5a).

**Fig. 5.**
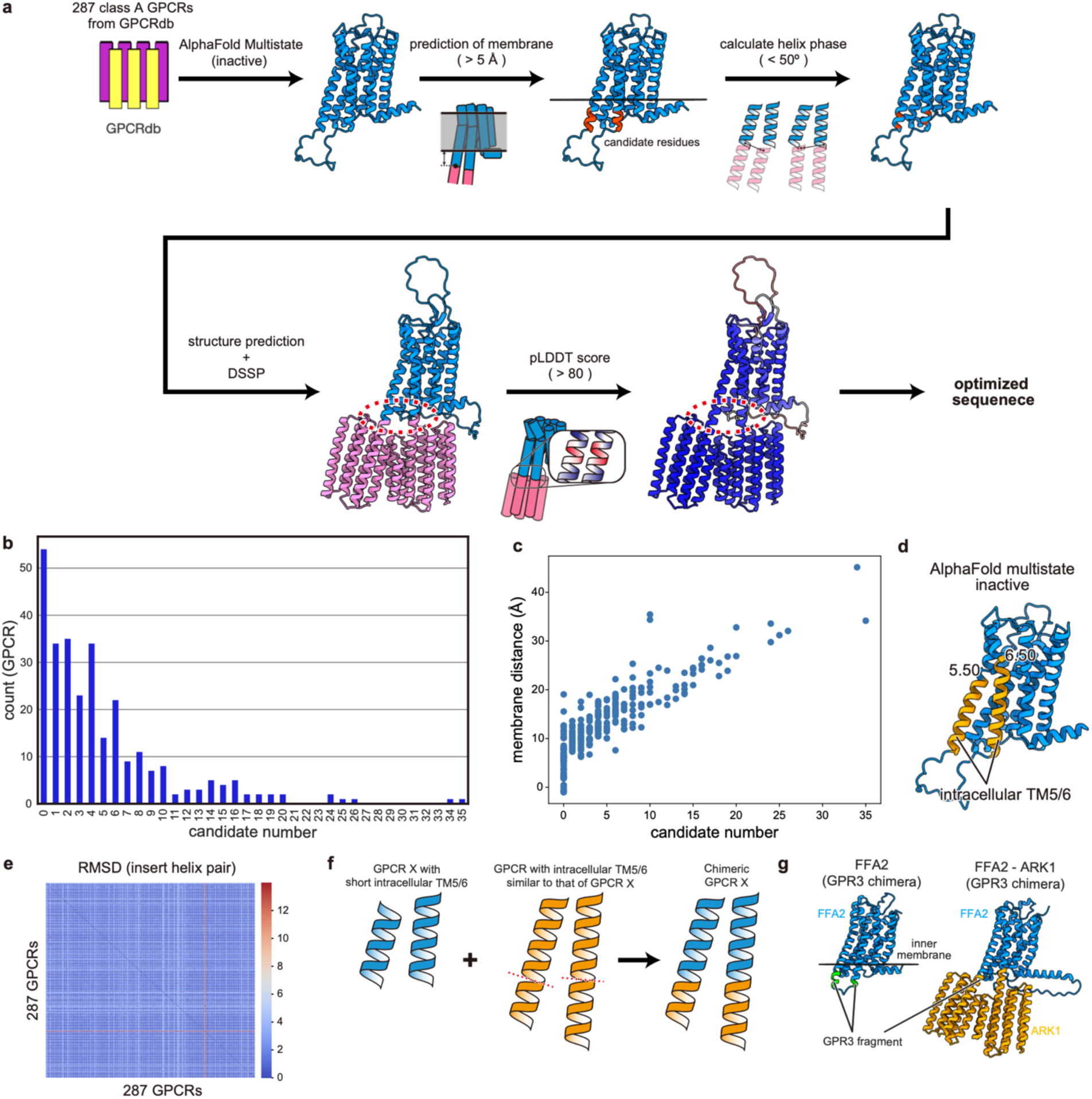
Comprehensive construct screening across the entire Class A GPCR. **a,** Automated screening workflow applied to all non-olfactory class A GPCRs (287 receptors) from GPCRdb. Inactive-state AlphaFold-Multistate models are first aligned with the predicted membrane plane, and residues whose Cα atoms lie > 5 Å outside the lipid bilayer are identified as candidate points for ARK1 fusion. For each candidate position on TM5 and TM6 suitable for ARK1 insertion, the helix phase between the two termini is calculated and candidates with Δθ < 50° are selected. Each potential construct is then rebuilt *in silico* and further filtered by DSSP and pLDDT score (> 80). **b,** Histogram depicting the number of acceptable TM5-TM6 fusion constructs identified per receptor. No viable candidates remained for 53 receptors. **c,** Correlation between the extent to which TM5 or TM6 protrudes beyond the membrane boundary in the wild-type model (y-axis, Å) and the number of acceptable constructs identified (x-axis) per receptor. **d,** Definition of insert helix pair. **e,** Heat-map of RMSD values for all 82,082 pairwise comparisons of insert helix pairs across the 287 receptors. **f,** Concept of chimeric extension strategy. A receptor (GPCR X, blue) with a short insert helix pair is grafted with a part of geometrically similar helix pair from a donor receptor (orange), generating a chimeric construct predicted to support a continuous TM5–TM6 fusion. **g,** Practical example using the free-fatty-acid receptor 2 (FFA2), whose native helix pair is too short for direct fusion. Insertion of the corresponding fragment from GPR3 yields a chimeric FFA2 suitable for ARK1 fusion.

This screen yielded at least one construct meeting all four criteria for over 200 receptors; however, 53 receptors were left with none (Fig. 5b). Further analysis revealed a strong correlation between the number of candidates constructs and the non-contact distance from the lipid bilayer to ARK1: receptors with intrinsically short TM5 and TM6 helices failed to meet the required threshold, causing all their designs to be filtered out (Fig. 5c).

To overcome this limitation, we refined our strategy by first computing pairwise structural homology of the TM5/TM6 intracellular segments across all 287 models (Fig. 5d,e). For each of the 53 outliers, we selected a donor GPCR with longer TM5/TM6 regions and the highest homology, and grafted its surplus intracellular helices onto the target receptor (Fig. 5f,g). Re-screening these extended constructs against our four criteria produced at least one viable ARK1-fusion design for every previously intractable GPCR.

### Determination of the antagonist-bound LPA2-ARK1 structure

Finally, we sought to validate the capability of the NOAH-ARK1 combination to determine the cryo-EM structure of a receptor using selected constructs. As a proof of concept, we chose lysophosphatidic acid receptor 2 (LPA2), which had not previously been structurally characterized in either its inactive or active conformation. A single selected LPA2-ARK1 fusion construct was expressed in HEK293 cells, then solubilized and purified in the presence of the widely used antagonist Ki16425 (Extended Data Fig. 9a,b). The sample was imaged using a Titan Krios microscope, yielding a high-resolution cryo-EM map of the Ki16425-bound LPA2-ARK1 complex at 2.99 Å resolution (Fig. 6a and Extended Data Fig. 9c-e). Notably, this map, like that of V2R-ARK1, exhibited high local resolution across the entire map, allowing modeling of nearly the entire receptor and ligand. Furthermore, the densities of several water molecules and a putative Na⁺ ion in the Na⁺ binding pocket were clearly visible (Fig. 6b,c and Extended Data Fig. 9f,g).

**Fig. 6.**
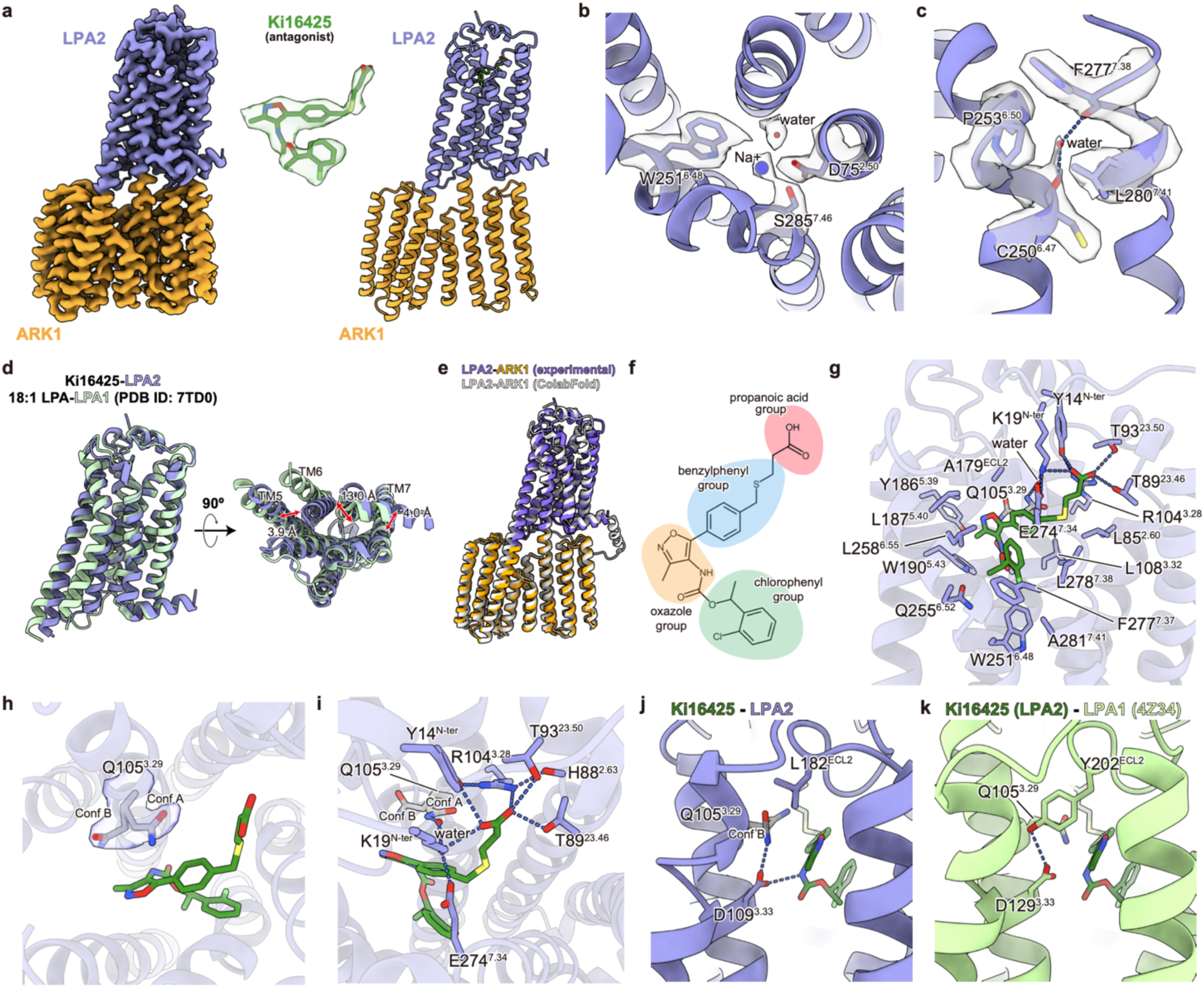
Structural analysis of LPA2-ARK1 bound to the antagonist Ki16425. **a,** Cryo-EM density map (left) and the corresponding atomic model (right) of the Ki16425-bound LPA2-ARK1 complex; the inset (center) overlays the ligand density with the atomic model. **b, c,** Close-up views of the conserved Na^+^ (b) and water molecule (c) within LPA2, features conserved across inactive-state class A GPCRs. The maps and models of water, ion, and surrounding residues are superposed. **d,** Superposition of inactive Ki16425-bound LPA2 with active 18:1-LPA-bound LPA1 (PDB: 7TD0), viewed parallel to the membrane (left) and from the intracellular side (right). **e,** Overlay of the experimentally determined LPA2-ARK1 structure with the ColabFold-predicted inactive LPA2 structure. **f,** Chemical structure of Ki16425 with pharmacophoric elements: propanoic acid, benzylphenyl, oxazole, and chloro-phenyl. **g,** Detailed ligand-binding mode of Ki16425. Hydrogen bonds are indicated by dashed lines. **h,** Two alternative conformations (conf. A and B) of Q105^3.29^. The cryo-EM density map is overlayed onto the atomic model of Q105^3.29^. **i,** Water-mediated hydrogen-bond network around the propanoic acid group of Ki16425. Hydrogen bonds are indicated by dashed lines. **j, k,** Hydrogen-bond network around Q^3.29^ and D^3.33^. Q105^3.29^ in conf. B interacts with D109^3.33^ in LPA2 (**j**), while this interaction is prevented by Y202^ECL2^ in LPA1 (**k**).

When comparing the LPA2-ARK1 structure with the previously reported active-state LPA1 structure, we observed an inward displacement of TM6 in LPA2-ARK1, a hallmark of the inactive state (Fig. 6d). Additionally, the structure closely matched the ColabFold-predicted model of inactive LPA2 (Fig. 6e). A detailed analysis of the ligand-binding site revealed that the propanoic acid group of Ki16425 forms extensive hydrogen bonds with the N-terminal region and ECL2 of LPA2, while the benzylphenyl, chlorophenyl, and oxazole groups interact extensively with amino acid residues at the orthosteric binding site across TMs 2, 3, 5, 6, and 7 (Fig. 6f,g).

Although Ki16425 functions as an antagonist for all three LPA receptors (LPA1, LPA2, and LPA3), it is known to have approximately 10-fold lower affinity for LPA2 compared to LPA1 or LPA3^39^. Sequence alignment revealed that among the residues interacting with Ki16425 in LPA2, only Q255^6.52^ is unique to LPA2 and is not conserved in LPA1 or LPA3 (Extended Data Fig. 10a). Interestingly, Q255^6.52^ in LPA2 would be positioned even closer to Ki16425 and form stronger interactions with the ligand than the equivalent residues, L275^6.52^ in LPA1 or L256^6.52^ in LPA3 (Extended Data Fig. 10b,c). Thus, it is unlikely that this difference alone would explain the lower affinity of Ki16425 for LPA2.

Further analysis identified a key feature potentially accounting for the affinity differences. The conserved residue Q105^3.29^, shared among LPA1-3, adopts two distinct conformations (conformer A and conformer B) in the LPA2 structure (Fig. 6h). In conformer A, Q105^3.29^ adopts a conformation similar to Q125^3.29^ in LPA1^40^, forming an NH-π interaction with the benzylphenyl group of Ki16425, and hydrogen bonds with Ki16425’s propanoic acid group mediated by water molecules and K19^ECL2^ (Fig. 6g-i). In contrast, in conformer B, Q105^3.29^ forms hydrogen bonds with D109^3.33^ but does not strongly interact with Ki16425 (Fig. 6j). Intriguingly, in LPA1, D129^3.33^ forms a hydrogen bond with Y202^ECL2^, preventing Q125^3.29^ from adopting conformer B due to steric clashes with Y202^ECL2^ (Fig. 6k, Extended Data Fig. 10d). Notably, in LPA2, the equivalent residue, L182^ECL2^, replaces the conserved Y202^ECL2^ found in LPA1 and LPA3. This suggests that the interaction between D^3.33^ and Y^ECL2^, observed only in LPA1 and LPA3, plays a crucial role in determining the conformation of Q105^3.29^. Therefore, the mixed conformations of Q105^3.29^ in LPA2 likely contribute to its reduced affinity for Ki16425 compared to LPA1 and LPA3, in which Q105^3.29^ exclusively adopts conformer A (Extended Data Fig. 10e).

In conclusion, the combination of ARK1 and NOAH facilitates the design of GPCR constructs suitable for high-resolution cryo-EM analysis, enabling visualization of ligand binding, water molecules, ions, and alternative side-chain conformations. This approach is applicable to other class A GPCRs, including LPA2, demonstrating the broad utility of this method.

## Discussion

G protein-coupled receptors (GPCRs) are integral to virtually every aspect of human physiology and represent a major class of drug targets. Among approved GPCR-directed drugs, approximately 44% act as antagonists and 55% as agonists^41^, underscoring the need for structural insights into both agonist- and antagonist-bound forms to deepen our molecular understanding and guide drug development.

In this study, we developed a universal pipeline that combines the *in silico* fusion-construct screening program NOAH with a *de novo*-designed fusion protein, ARK1, to facilitate GPCR structure determination. We validated this approach by solving the structures of three class A GPCRs in their antagonist- or agonist-bound states: tolvaptan-bound V2R, OPC5-bound V2R, and icatibant-bound B2R, revealing new facets of their activation and deactivation mechanisms. We further demonstrated the robustness of NOAH-ARK1 by determining the tolvaptan-bound V2R structure at higher resolution and extending its applicability to the antagonist-bound LPA2 receptor. Among the four structures we report, three (OPC5-bound V2R, icatibant-bound B2R, and Ki16425-bound LPA2) were previously undescribed. Although a tolvaptan-bound V2R structure was recently determined using a similar BRIL-Fab fusion strategy^42^, our tolvaptan-bound V2R-BRIL structure achieved notably higher receptor resolution, particularly in the extracellular binding pocket, and was surpassed further by our tolvaptan-bound V2R-ARK1 structure. These findings highlight the importance of carefully optimizing the linker between the receptor and fusion partner using multiple criteria to achieve higher-resolution data.

While this work was in progress, several additional methods for GPCR structure determination emerged, including the KOR ICL3 graft with nanobody addition^13^, the PGS fusion approach^14^, the calcineurin fusion technique^15^, and a “glue complex” strategy that combines BRIL, an ALFA tag, an anti-BRIL Fab, and a dual nanobody-fusion system^16^. However, each of these approaches requires extensive experimental screening to identify suitable linker sequences. Although ColabFold or AlphaFold2 have occasionally been employed to optimize linker sequences^33^, our results indicate their predictions alone are often overly optimistic, necessitating additional criteria to effectively guide construct selection. By incorporating three additional criteria, our NOAH program addresses this challenge and significantly reduces the burden of trial-and-error screening.

Notably, the NOAH-ARK system not only eliminates the need for extensive experimental screening but also simplifies workflows by removing the requirement to prepare multiple proteins (e.g., nanobodies, Fab, or NbFab), as necessary in other methods such as BRIL-Fab, KOR’s ICL3, or glue complex approaches^13,16^. Moreover, NOAH-ARK enhances receptor expression levels and yields higher-resolution maps, especially around ligand-binding pockets, enabling accurate modeling of ligands and, in some cases, even water molecules and metal ions. Additionally, unlike the glue complex approach, which is difficult to apply to GPCRs lacking helix 8, the NOAH-ARK system can be applied to all non-olfactory class A GPCRs.

The concept of a *de novo*-designed fusion partner was also recently explored in a preprint describing “Clip” proteins generated via RFdiffusion^43^. However, creating an optimal fusion partner with sufficient size, asymmetry, and stability—essential for high-resolution structural analysis, including modeling water molecules and ions—remains challenging with RFdiffusion, as Clip proteins are only 17-19 kDa. Additionally, because each receptor requires a distinct Clip protein, further gene synthesis is necessary for each construct. In contrast, our BDP method circumvents these limitations by producing a versatile fusion partner that can be swapped among class A GPCRs, underscoring the broad applicability of the NOAH-ARK platform.

Although AlphaFold2 and ColabFold can predict GPCR structures and aid in evaluating fusion constructs, they cannot predict ligand binding. More recent software, such as AlphaFold3 (AF3), Boltz-1, and Chai-1, can, in principle, model protein-ligand complexes^44–46^. However, we found that AF3 predictions for OPC5- and icatibant-bound receptors deviated markedly from our experimentally determined structures (Extended Data Fig. 11a,b). This underscores the continued necessity of experimentally determining GPCR-ligand complexes to ensure reliable mechanistic insights and to support drug discovery efforts.

In conclusion, the method presented here offers a universal strategy for determining GPCR structures in various ligand-bound states, providing new opportunities to advance our understanding of GPCR structure and function, and to inform future drug design.

## Data Availability

The raw images of tolvaptan-bound V2R-BRIL, OPC51803-bound V2R-BRIL, icatibant-bound B2R-BRIL, tolvaptan-bound V2R-ARK1, Ki16425-bound LPA2-ARK1 before motion correction have been deposited in the Electron Microscopy Public Image Archive under accession EMPIAR-xxxxx. The cryo-EM density map and atomic coordinates for icatibant-bound B2R-BRIL (overall or TM), tolvaptan-bound V2R-BRIL (overall or TM), OPC51803-bound V2R-BRIL (TM), tolvaptan-bound V2R-ARK1, Ki16425-bound LPA2-ARK1 have been deposited in the Electron Microscopy DataBank: EMD-64536, EMD-64537, EMD-64538, EMD-64539, EMD-64540, EMD-64541 and EMD-64542, and PDB under accessions: 9UVT, 9UVU, 9UVV, 9UVW, 9UVX, 9UVY and 9UVZ, respectively. All other data are provided in this article or from the corresponding author on reasonable request.

## Acknowledgements

We thank T. Kusakizako and Y. Sakamaki (UTokyo) for help with cryo-EM data collection, K. Hasegawa for administrative support, and Otsuka Pharmaceutical Co., Ltd. for providing Tolvaptan and OPC51803. The electron microscopic observations were supported by Research Support Project for Life Science and Drug Discovery (Basis for Supporting Innovative Drug Discovery and Life Science Research [BINDS]) from the Japan Agency for Medical Research and Development (AMED) under grant number JP22ama121002, JP23ama121002, and JP24ama121002. We also acknowledge ChatGPT, a multimodal large language model created by OpenAI, for providing guidance to improve the readability of this manuscript. Note that, after using this tool, we reviewed and edited the content as needed and took full responsibility for the content of the publication.

This work was supported by the Uehiro Foundation (H.E.K.), AMED (24bm1123057h0001 to H.E.K.), JSPS KAKENHI (JP24KJ0981 to A.K, JP22H05426/JP24H01139 to N.K., JP23KJ0363/JP24K18286/JP25H01338 to K.Kawakami, JP24K18060/JP25H02243 to K.Kobayashi, JP24H02262/JP25K09525 to M.F., JP21H04969/JP22H00400/JP25H01338 to H.E.K.), JST PRESTO (JPMJPR24OF to M.F.), JST FOREST (JPMJFR204S to H.E.K.), JST CREST (JPMJCR21P3/JPMJCR23B1 to H.E.K.), 10th JACI Prize for Encouraging Young Researcher (N.K.), the Senri Life Science Foundation (K.Kawakami), the Uehara Memorial Foundation (K.Kawakami), The Naito Science & Engineering Foundation (K.Kawakami), and a research grant from Ono Pharmaceutical (H.E.K.).

## Author Contributions

A.K. and T.E.M. developed the NOAH program. A.K. and N.K. designed the ARK1 protein. A.K. performed the biochemical analysis. A.K. and K.U. performed the cryo-EM analysis with assistance from K.Kobayashi, T.E.M., and M.F. K.Kobayashi and Y.G. provided valuable input on structural considerations. A.K., K.Kawakami, and H.E.K. wrote the paper with help from all authors. H.E.K. supervised the research.

---

**Extended Data Fig. 1.**
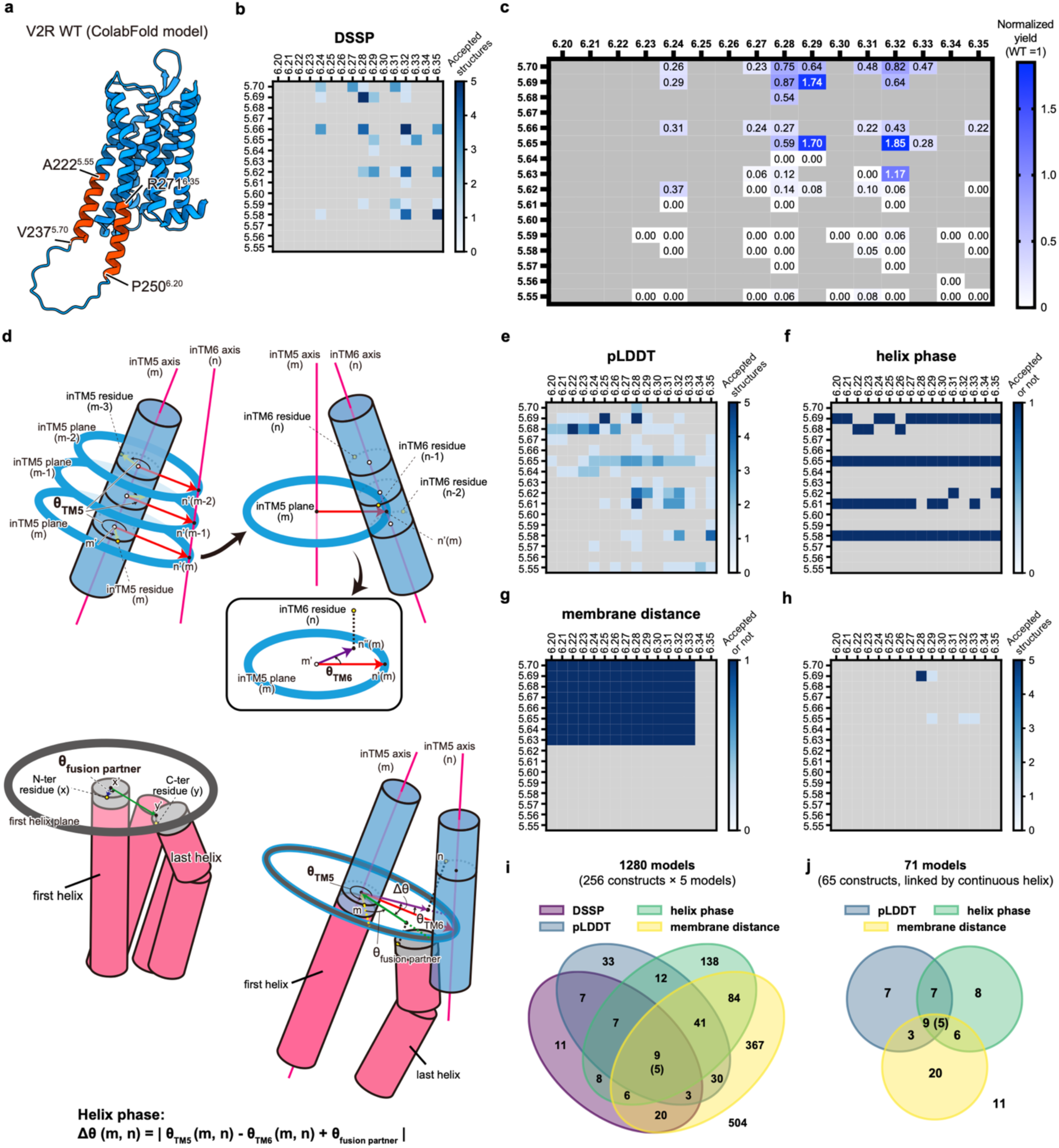
Screening criteria developed through the structural analysis of V2R. **a,** Candidate positions for BRIL insertion within the TM5 and TM6 region of the V2R. **b,** DSSP filter. Heatmap (blue scale, 0–5) indicating how many of the five ColabFold-predicted structures retain a continuous α-helix spanning both receptor and BRIL for each TM5/TM6 insertion pair. **c,** Expression screening. Normalized FSEC peak heights (WT = 1; grey → blue) for the 65 constructs passing the DSSP filter. **d,** Geometric definition of the “helix phase” (Δθ): schematic explaining how the relative angular registers of TM5, TM6 and the BRIL helices are measured. **e,** pLDDT filter. Heatmap showing the number of models (0–5) with a pLDDT score higher than 80 at the TM5-TM6 junction. **f,** Helix-phase filter. Binary map indicating constructs with Δθ smaller than 50° (dark blue = accepted). **g,** Membrane-distance filter. Binary map identifying constructs in which all BRIL residues lie more than 5 Å outside the PPM3^21^-predicted membrane boundary. **h,** Composite filter integrating all four criteria (DSSP, pLDDT, helix phase, and membrane boundary gap); only five constructs fulfill all criteria. **i,** Venn diagram summarizing overlaps among the four individual criteria across all 1,280 ColabFold models (256 constructs × 5 predictions). **j,** Venn diagram illustrating how three different filters (pLDDT, helix phase, and membrane distance) narrow the candidates to 9 models (corresponding to 5 constructs) out of the 71 models (65 constructs) that initially passed the DSSP filter.

**Extended Data Fig. 2.**
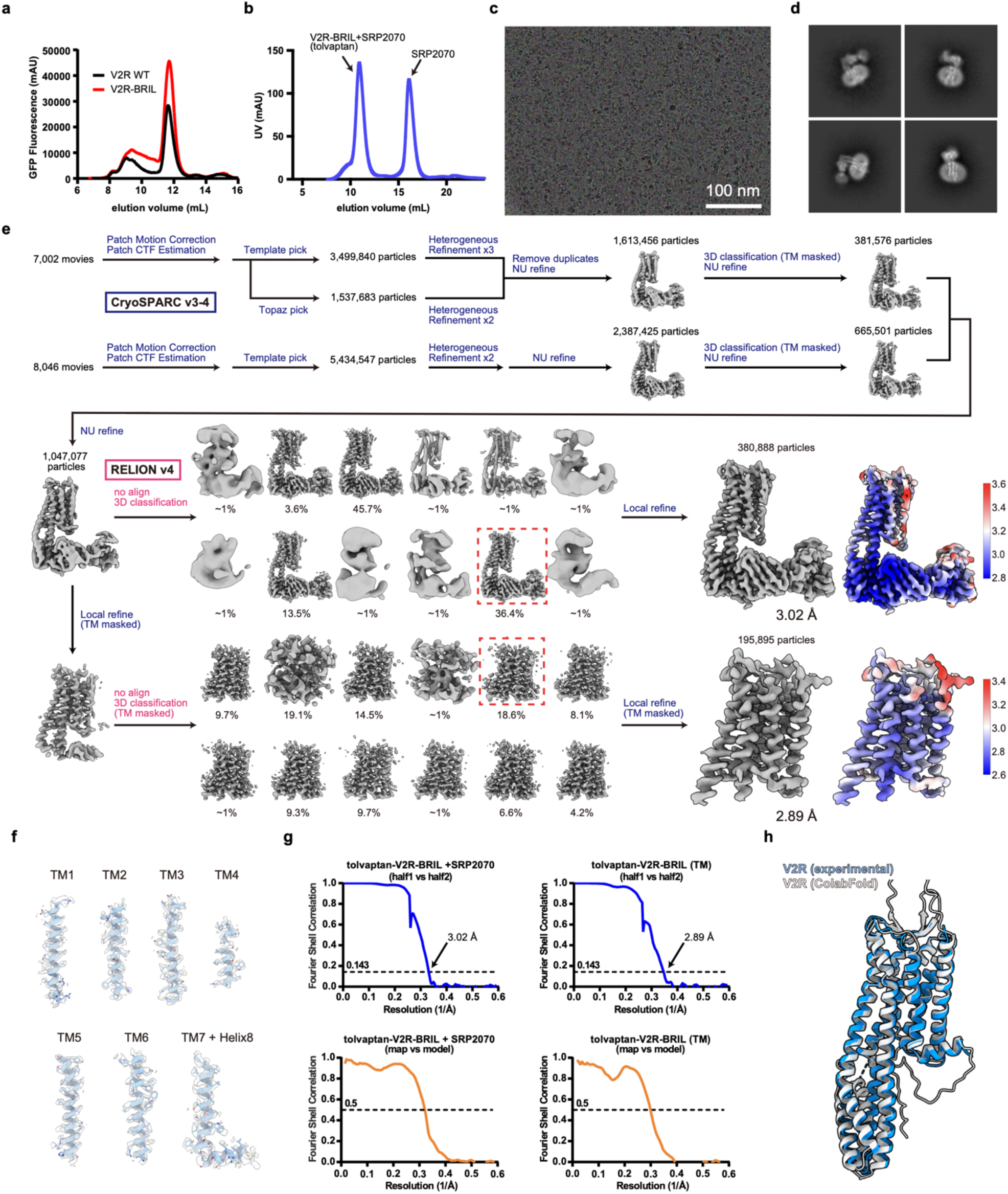
Cryo-EM analysis of tolvaptan-bound V2R-BRIL SRP2070 Fab complex. **a,** Representative FSEC traces of WT (black) or BRIL-fused (red) V2R-GFP constructs. **b,** Representative SEC trace of the purified V2R-BRIL complex with the SRP2070 Fab antibody. **c,** Representative cryo-EM micrograph. **d,** Representative 2D class averages. **e,** Data-processing workflow with the final cryo-EM maps colored by local resolution. **f,** Superposition of the cryo-EM density map and the atomic model for TM helices. **g,** Fourier shell correlation (FSC) between the two independently refined half maps (top left: overall structure, top right: TM regions) and between the model and the map calculated for the model refined against the full reconstruction (bottom left: overall structure, bottom right: TM regions). **h,** Comparison between the experimentally determined cryo-EM structure (blue) and ColabFold-predicted structure (white).

**Extended Data Fig. 3.**
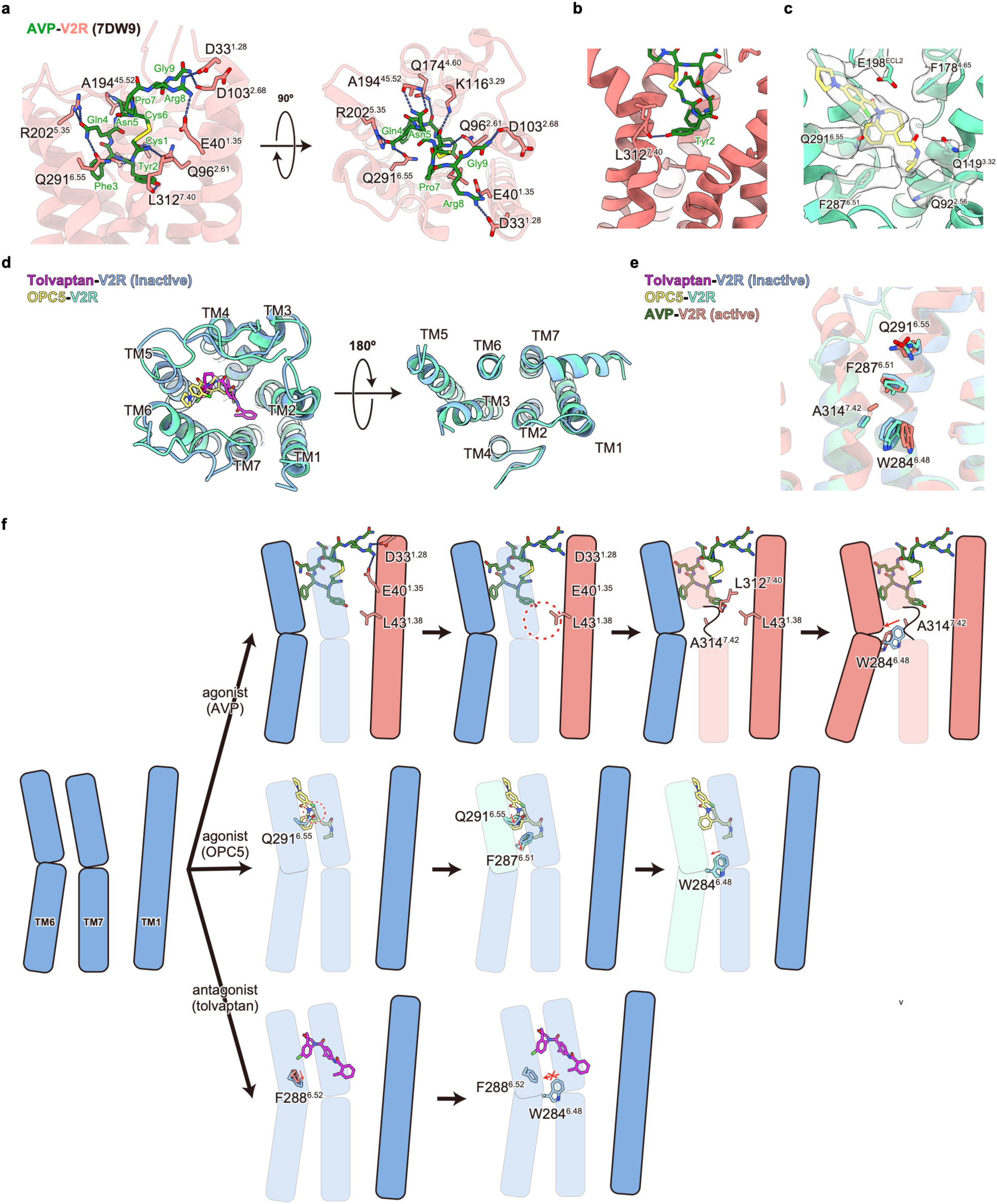
Ligand recognition and receptor activation/deactivation mechanism of V2R. **a,** Binding mode of AVP in the active-state V2R structure (PDB: 7DW9), viewed parallel to the membrane (left) and from the extracellular side (right). Dashed lines indicate hydrogen bonds and salt bridges. **b,** Close-up view of the hydrogen bond formed between AVP Tyr2 and the backbone carbonyl of L312^7.40^. **c,** Cryo-EM density overlaid on the atomic model of the partial agonist OPC5 bound to V2R. **d,** Structural comparison of the inactive, tolvaptan-bound V2R and the OPC5-bound state, viewed from the extracellular side (left) and intracellular side (right). **e,** Structural superposition around the conserved toggle-switch residue. **f,** Schematic model of V2R regulation by the full agonist AVP, partial agonist OPC5, and antagonist tolvaptan. First, AVP pulls TM1 inward via salt bridges involving Arg8, and inwardly shifted TM1 presses on TM7 via L43^1.38^. This induces partial unwinding and inward displacement of TM7, stabilized by a hydrogen bond between AVP Tyr2 and L312^7.40^. The repositioned TM7 then interacts with the toggle-switch W284^6.48^, triggering a kink around P286^6.50^ and outward movement of TM6, thus establishing a fully active conformation (top). Second, OPC5 interacts with a TM6 residue (Q291^6.55^), inducing a subtle rotation and displacement of TM6 and the toggle-switch residue. Therefore, OPC5 activates V2R by directly promoting TM6 movement, which is distinct from AVP-induced activation (middle). Third, tolvaptan shares a backbone scaffold with OPC5, however, it binds the receptor in a ∼180° rotated orientation. Tolvaptan binds the F288^6.52^ residue, sterically preventing both the inward shift of TM7 and the outward rotation of TM6. This locks the receptor in its inactive state (bottom).

**Extended Data Fig. 4.**
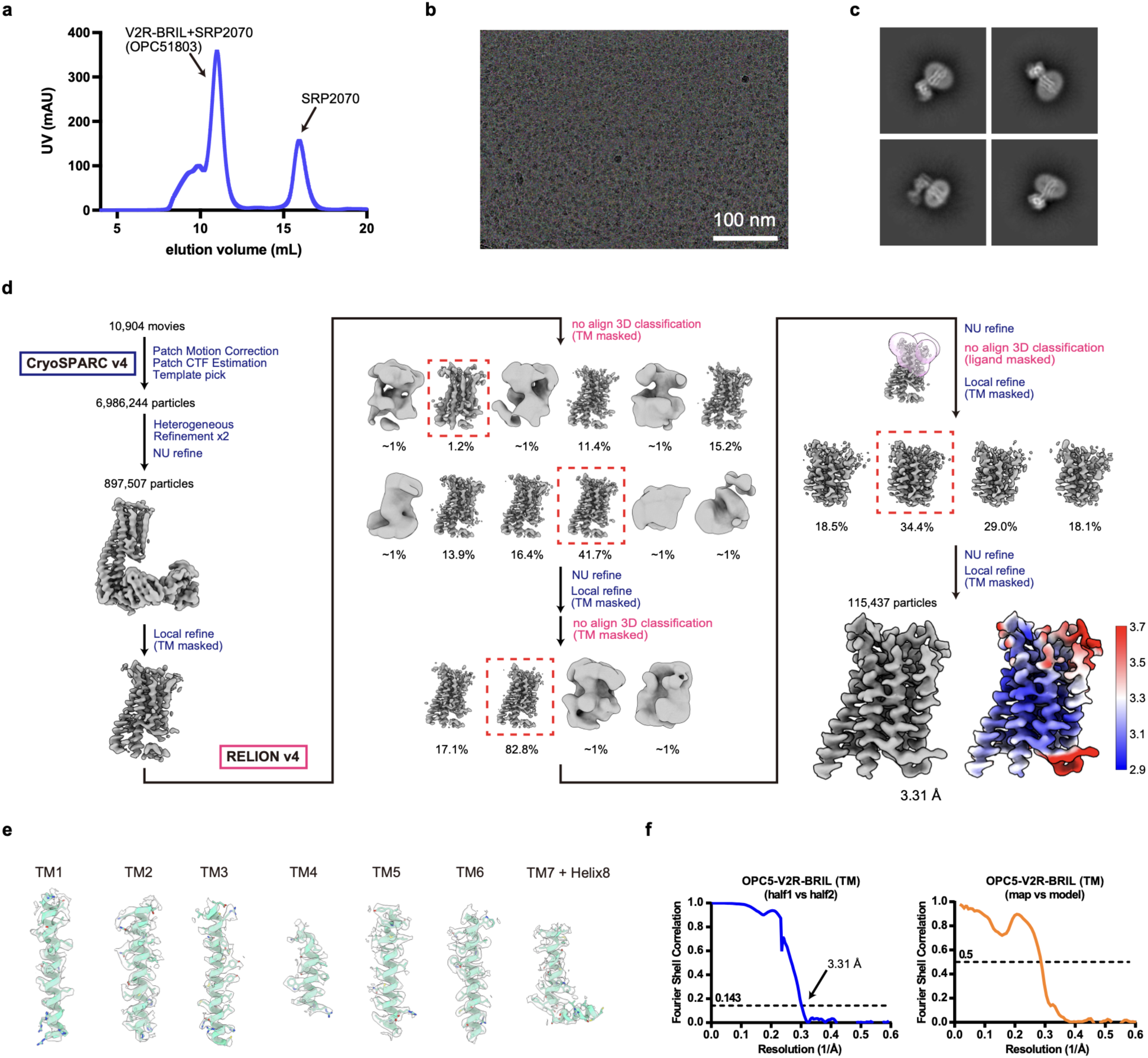
Cryo-EM analysis of OPC51803-bound V2R-BRIL SRP2070 Fab complex. **a,** Representative SEC trace. **b,** Representative cryo-EM micrograph. **c,** Representative 2D class averages. **d,** Data processing workflow with the final cryo-EM map colored by local resolution. **e,** Superposition of the cryo-EM density map and the atomic model for TM helices. **f,** FSC between the two independently refined half maps (left) and between the model and the map calculated for the model refined against the full reconstruction (right).

**Extended Data Fig. 5.**
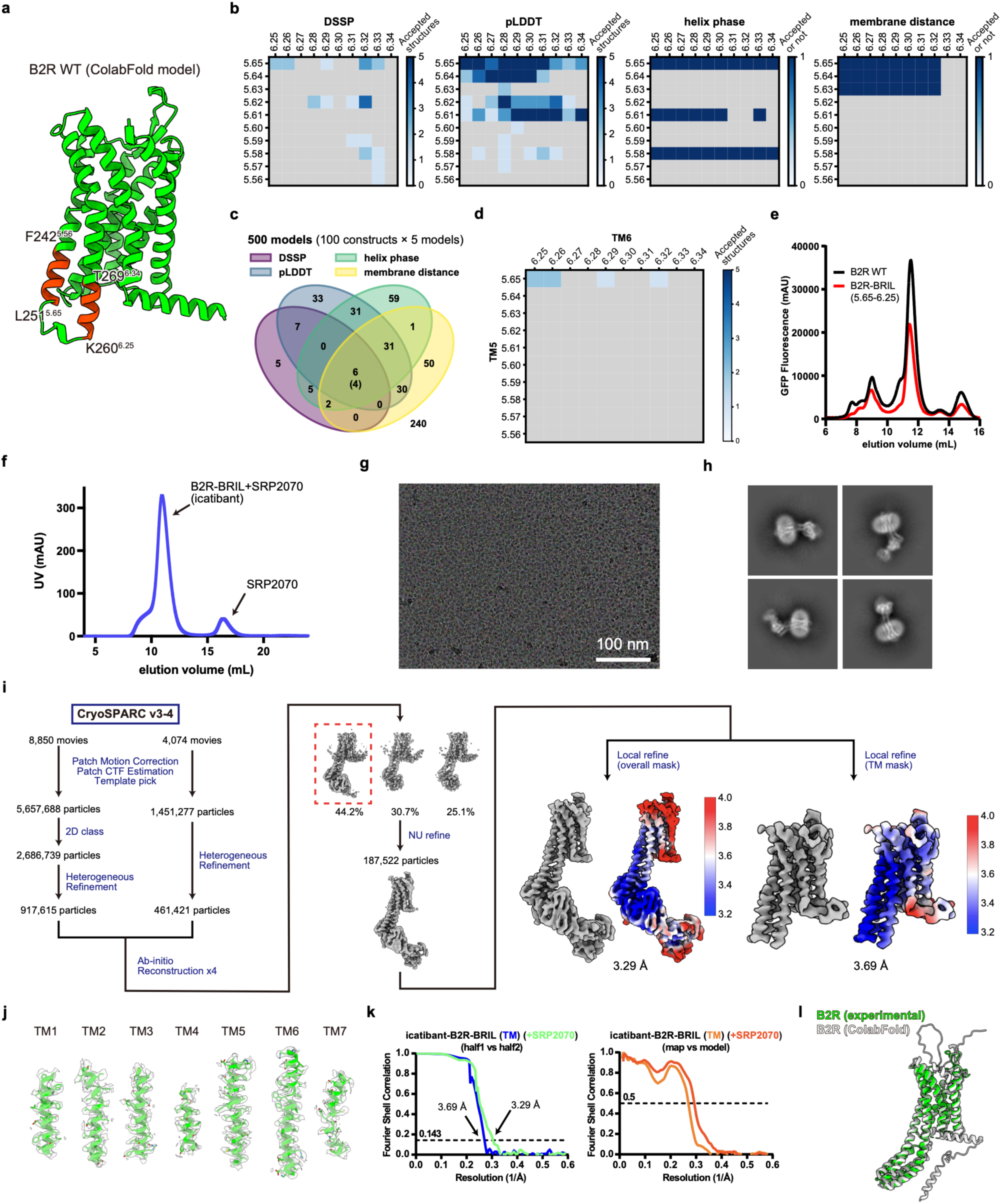
Cryo-EM strucutre of the icatibant-bound B2R-BRIL–anti-BRIL Fab complex. **a,** Candidate positions for BRIL insertion within the TM5 and TM6 region of B2R. **b,** *In-silico* selection of 100 TM5–TM6 fusion constructs (five ColabFold models each). Constructs were selected using DSSP, pLDDT, helix phase, and membrane boundary gap filters. **c,** Venn diagram showing how the four individual filters narrow the candidates from 500 predicted models. Number in parentheses represents the number of constructs passing all four filters. **d,** Composite filter integrating all four criteria. **e,** Representative FSEC traces of WT (black) or BRIL-fused (red) B2R- GFP constructs. **f,** Representative SEC trace of the purified icatibant-bound B2R-BRIL complex with the SRP2070 Fab antibody. **g,** Representative cryo-EM micrograph. **h,** Representative 2D class averages. **i,** Data processing workflow with the final cryo-EM maps colored by local resolution. **j,** Superposition of the cryo-EM density map and the atomic model for TM helices. **k,** FSC between the two independently refined half maps (TM regions in blue, overall structure in light green) and between the model and the map calculated for the model refined against the full reconstruction (TM regions in orange, overall structure in light orange). **l,** Comparison between the experimentally determined cryo-EM structure and ColabFold-predicted structure.

**Extended Data Fig. 6.**
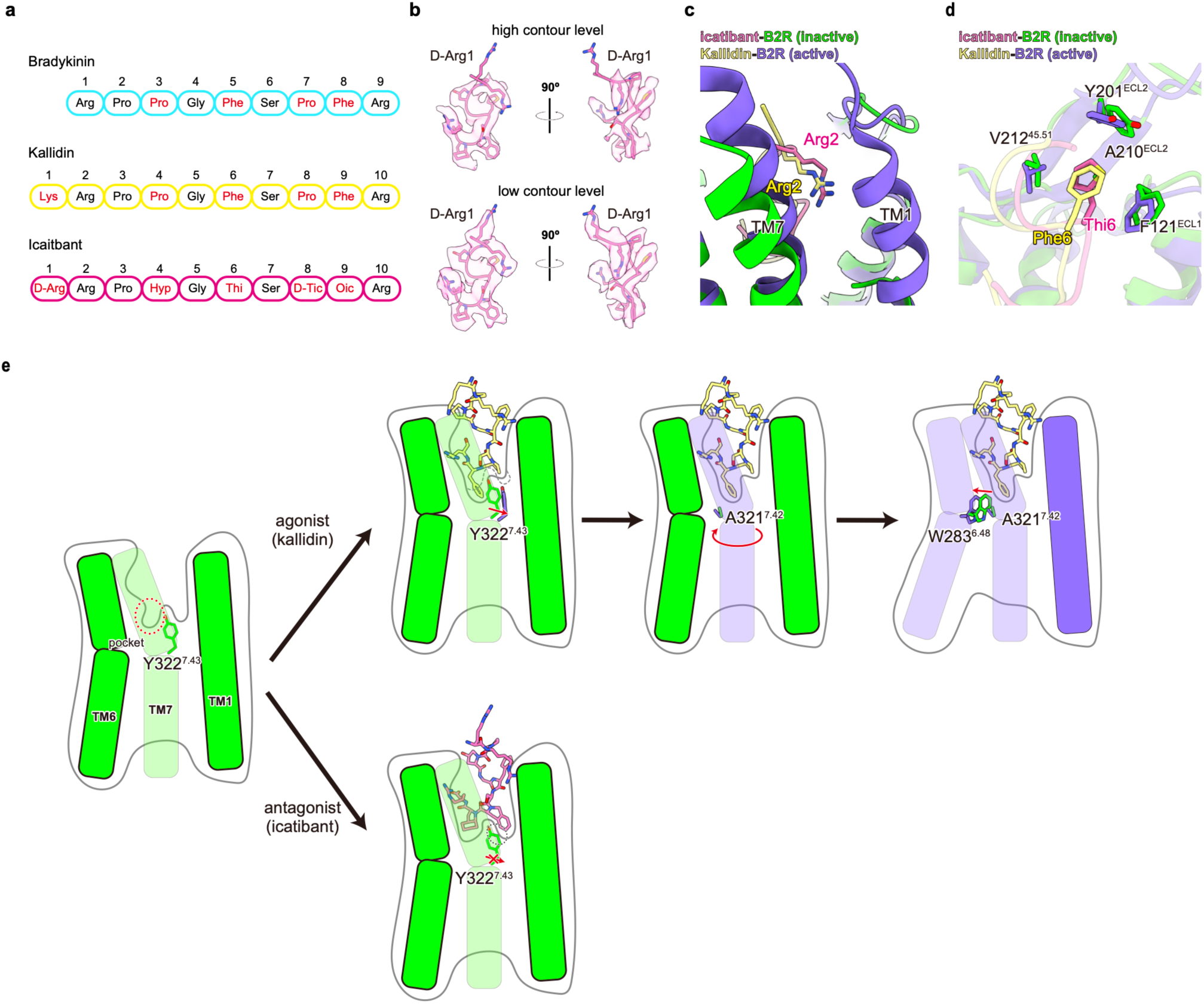
Ligand recognition and receptor activation/deactivation mechanism of B2R. **a,** Amino-acid sequences of bradykinin, kallidin, and icatibant. **b,** Cryo-EM density of icatibant contoured at high (upper) and low (lower) threshold, shown together with the corresponding atomic model. Note that the density of D-Arg1 is weaker than for other residues, suggesting that D-Arg1 does not strongly interact with the receptor. **c,d,** Binding mode of Arg2/Arg2 (c), and Phe6/Thi6 (d) in kallidin and icatibant, respectively. **e,** Schematic model of B2R regulation by the agonist kallidin and antagonist icatibant. Kallidin or bradykinin inserts the C-terminal residue, Phe9, deeply into the binding pocket, pushing Y322^7.43^ downward. The downward movement of Y322^7.43^ induces clockwise rotation of TM7, causing a steric clash between A321^7.42^ and W283^6.48^. This clash triggers outward movement of TM6, establishing the active receptor conformation (top). In contrast, icatibant, with bulkier D-Tic8 and Oic9 residues, does not induce the downward shift of Y322^7.43^ but instead sterically blocks its movement, preventing activation (bottom).

**Extended Data Fig. 7.**
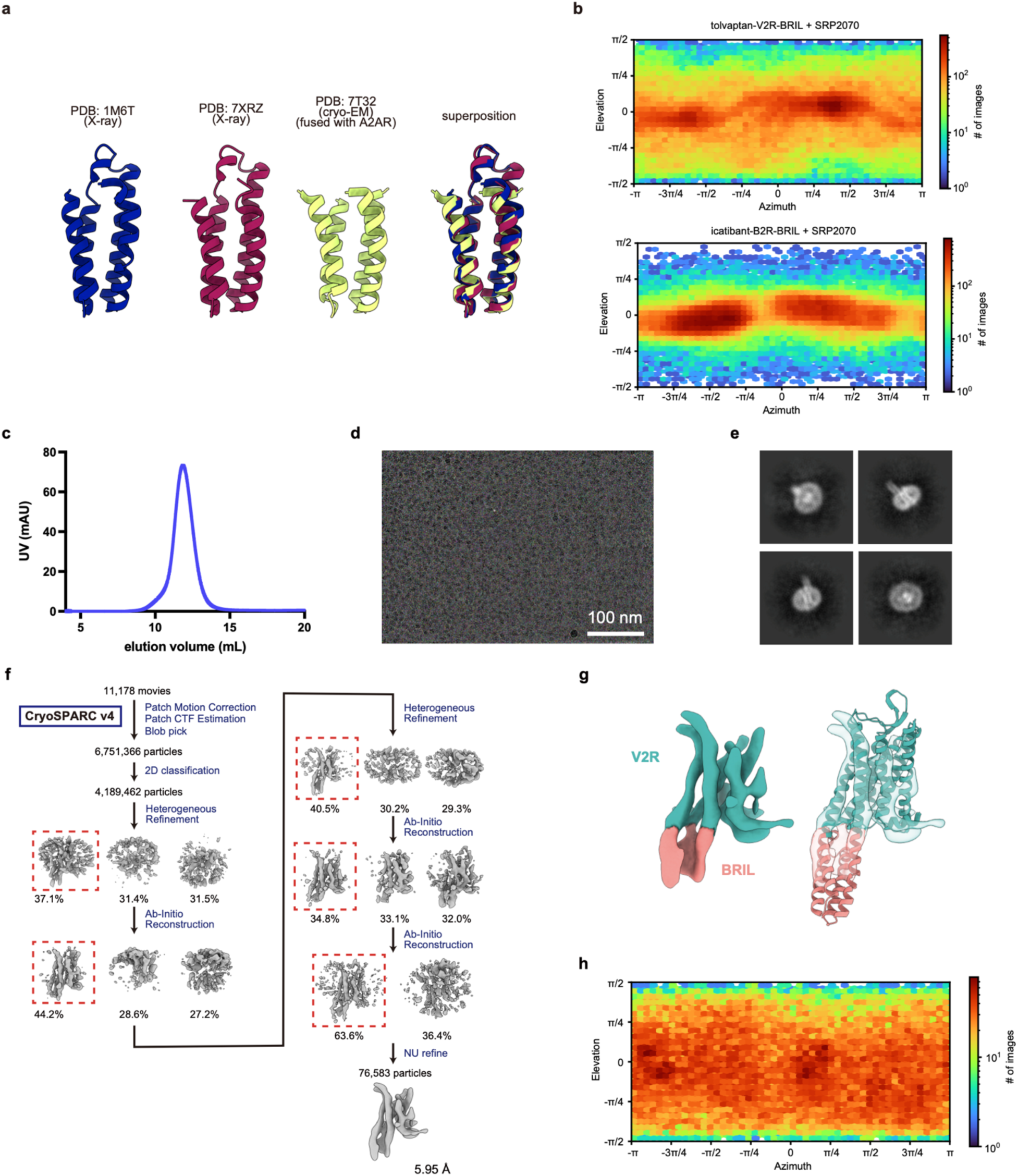
Impact of BRIL and Fab on structural analysis. **a,** Comparison of BRIL structures fused to various receptors, extracted from receptor structures deposited in the PDB. The C-terminal end of BRIL exhibits considerable flexibility. **b,** Orientation distribution plots of the tolvaptan-bound V2R-BRIL-SRP2070 Fab complex (top) and icatibant-bound B2R-BRIL-SRP2070 Fab complex (bottom). **c,** Representative SEC trace. **d,** Representative cryo-EM micrograph. **e,** Representative 2D class averages. **f,** Data processing workflow. **g,** Cryo-EM density map colored by V2R in green and BRIL in salmon (left), and overlay of the density map and model of the tolvaptan-bound V2R-BRIL without Fab antibody (right). **h,** Orientation distribution plot of tolvaptan-bound V2R-BRIL, suggesting that while Fab antibody is necessary for high-resolution cryo-EM analysis, it introduces an undesired orientation bias on the cryo-EM grids.

**Extended Data Fig. 8.**
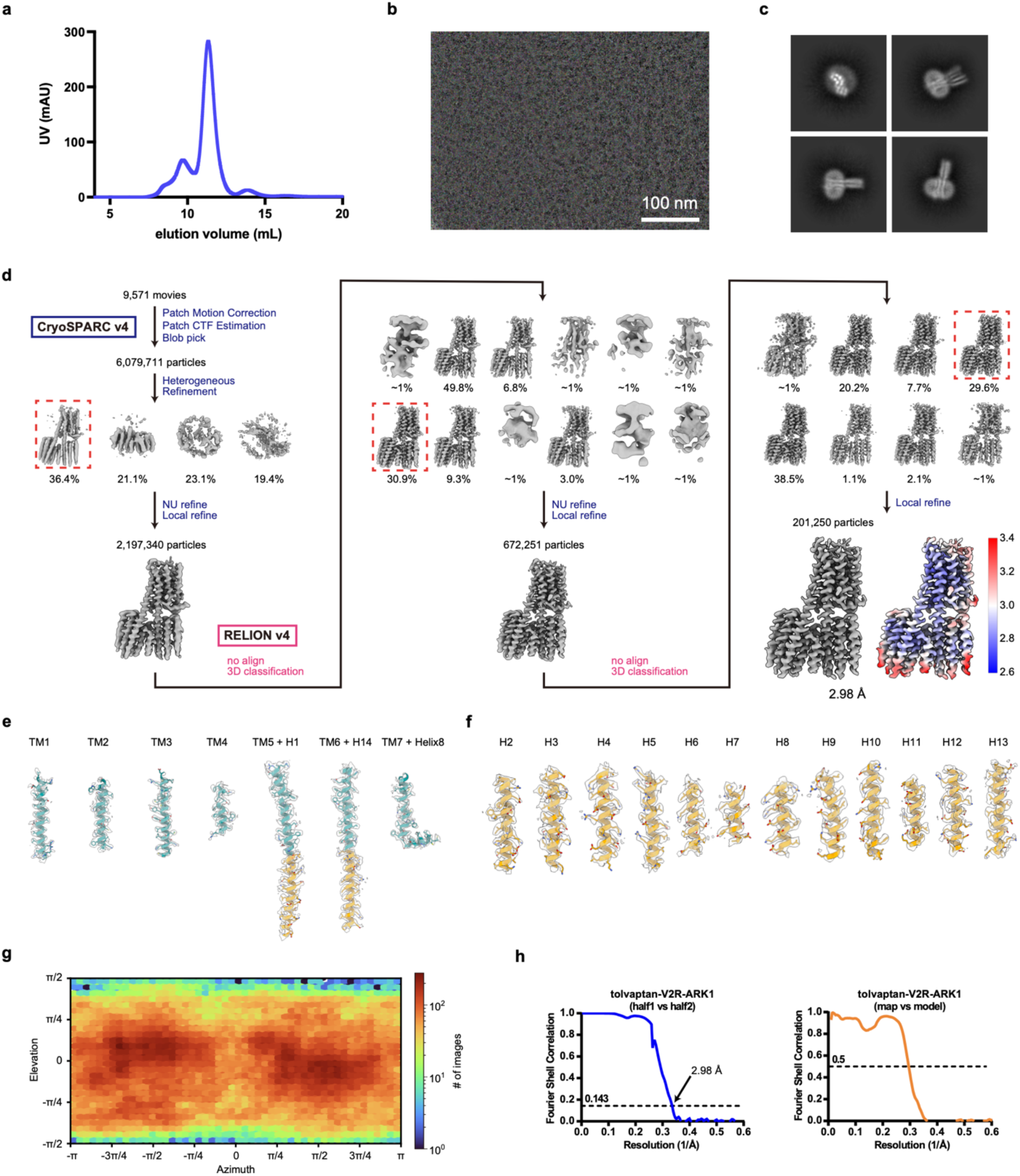
Cryo-EM analysis of tolvaptan-bound V2R-ARK1. **a,** Representative SEC trace. **b,** Representative cryo-EM micrograph. **c,** Representative 2D class averages. **d,** Data processing workflow with the final cryo-EM map colored by local resolution. **e,** Superposition of the cryo-EM density map and the atomic model for TM helices of the V2R region. **f,** Superposition of the cryo-EM density map and the atomic model for 13 helices of the ARK1 region. **g,** Orientation distribution plot of the tolvaptan-bound V2R-ARK1. Note that orientation bias is reduced compared to that observed for the V2R-BRIL-SRP2070 Fab complex. **h,** FSC between the two independently refined half maps (left) and between the model and the map calculated for the model refined against the full reconstruction (right).

**Extended Data Fig. 9.**
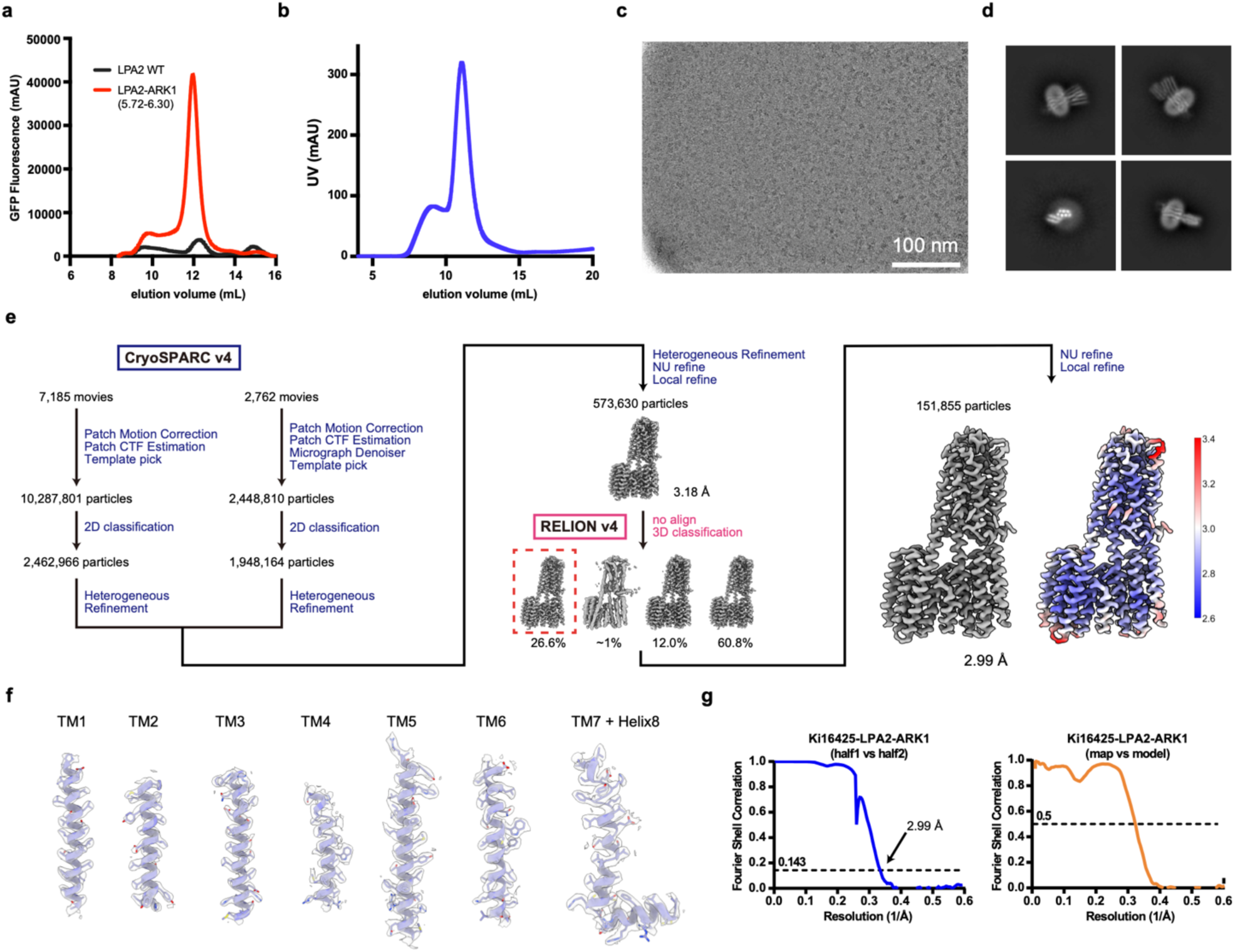
Cryo-EM analysis of Ki16425-bound LPA2-ARK1. **a,** Representative FSEC traces of WT or ARK1-fused LPA2-GFP constructs. **b,** Representative SEC trace. **c,** Representative cryo-EM micrograph. **d,** Representative 2D class averages. **e,** Data processing workflow with the final cryo-EM map colored by local resolution. **f,** Superposition of the cryo-EM density map and the atomic model for TM helices of the LPA2 region. **g,** FSC between the two independently refined half maps (left) and between the model and the map calculated for the model refined against the full reconstruction (right).

**Extended Data Fig. 10.**
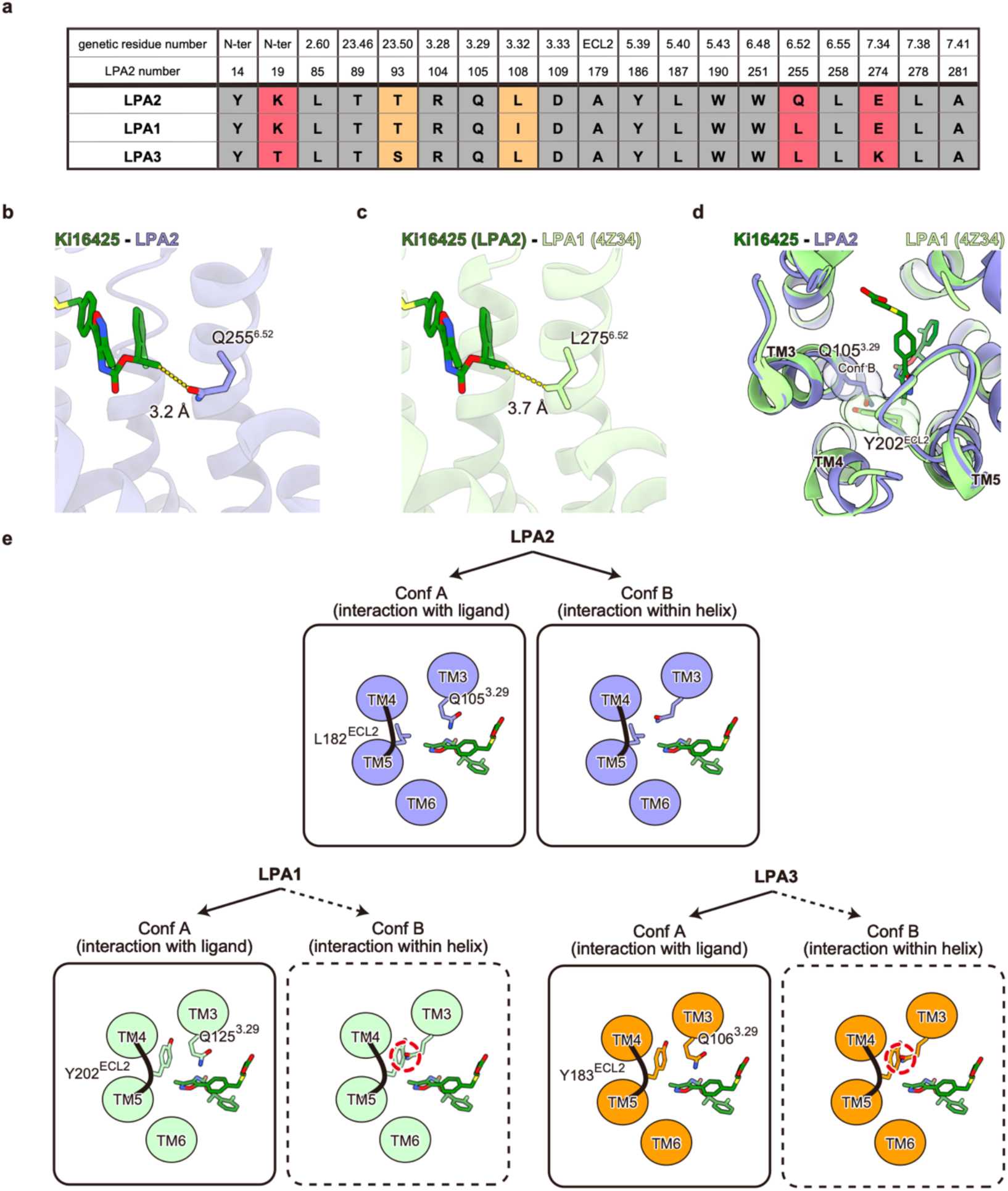
Structural insights for the lower affinity of Ki16425 to LPA2 compared to LPA1 and LPA3. **a,** Amino acid sequence alignment highlighting residues that interact with Ki16425 in the LPA2-ARK1 structure among LPA1-LPA3. Conserved residues are shaded grey, structurally similar residues are highlighted in orange, and notably distinct residues are indicated in red. **b,** Close-up view of the interaction between Q255^6.52^ and Ki16425 in LPA2. **c,** Model of Ki16425 docked into LPA1. The distance between Ki16425 and L275^6.52^ in LPA1 is longer than the equivalent distance between Ki16425 and Q255^6.52^ in the LPA2 structure. **d,** Structural overlay of Ki16425-bound LPA2 and antagonist-bound LPA1, highlighting a steric clash between Ki16425 and Y202^ECL2^ in LPA1. **e,** Schematic representation of why Ki16425 shows weaker affinity for LPA2 compared to LPA1 and LPA3. In LPA2, Gln105^3.29^, a putative key residue for Ki16425 binding, adopts two conformations. In conformer A, Gln105^3.29^ strongly interacts with Ki16425, stabilizing tis binding. However, in conformer B, Gln105^3.29^ flips inward, forming an intrahelical hydrogen bond with D109^3.33^, thus losing contact with the ligand (top). In contrast, LPA1 and LPA3 possess a tyrosine residue in ECL2 (Y202^ECL2^ in LPA1 and Y183 ^ECL2^ in LPA3) that restricts Q^3.29^ to conformer A, maintaining a stable interaction with Ki16425 and consequently higher binding affinity (bottom).

**Extended Data Fig. 11.**
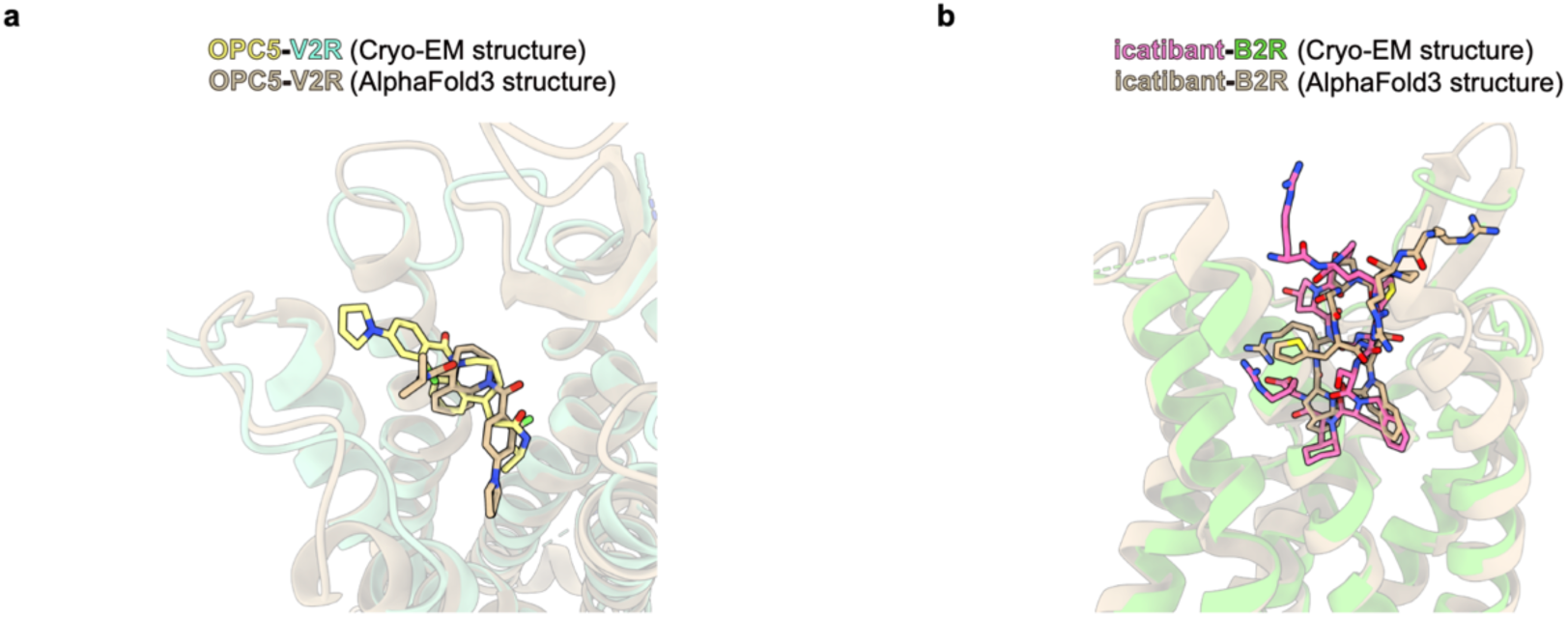
Comparison between cryo-EM and AlphaFold3-predicted structures. **a,** Structural comparison between the experimentally determined and AlphaFold3-predicted OPC5-bound V2R structures. **b,** Structural comparison between experimentally determined and AlphaFold3-predicted icatibant-bound B2R structure.

