## Supplementary Table 1 for "A Rapid and Universal Pipeline for High-Resolution GPCR Structure Determination through *In Silico* Construct Optimization and de novo Protein Design"

### Cryo-EM data collection, refinement and validation statistics

| | V2R (Tolvaptan)-BRIL+SRP2070Fab | | V2R (OPC51803)-BRIL+SRP2070 Fab |
|---|---|---|---|
| | Overall (EMDB-64538) (PDB 9UVV) | TM focused (EMDB-64539) (PDB 9UVW) | TM focused (EMDB-64540) (PDB 9UVX) |
| **Data collection and processing** | | | |
| Magnification | 105,000 | | 105,000 |
| Voltage (kV) | 300 | | 300 |
| Electron exposure (e–/Å$^2$) | 50.0 | | 45.6 |
| Defocus range (μm) | -0.6 to -1.8 | | -0.6 to -1.8 |
| Pixel size (Å) | 0.83 | | 0.83 |
| Symmetry imposed | C1 | C1 | C1 |
| Initial particle images (no.) | 10,472,070 | | 6,986,244 |
| Final particle images (no.) | 380,888 | 195,895 | 115,437 |
| Map resolution (Å) | 3.02 | 2.89 | 3.31 |
| FSC threshold | 0.143 | 0.143 | 0.143 |
| **Refinement** | | | |
| Initial model used (PDB code) | AF2 model, 7XRZ for Fab | AF2 model | AF2 model |
| Map sharpening *B* factor (Å$^2$) | 97.1 | 77.4 | 95.2 |
| Model composition | | | |
| Non-hydrogen atoms | 6,121 | 2,113 | 2,089 |
| Protein residues | 785 | 266 | 263 |
| Ligands | 1 | 1 | 1 |
| *B* factors (Å$^2$) | | | |
| Protein | 108.765 | 98.009 | 99.121 |
| Ligand | 66.269 | 65.540 | 94.401 |
| R.m.s. deviations | | | |
| Bond lengths (Å) | 0.009 | 0.008 | 0.009 |
| Bond angles (°) | 1.64 | 1.54 | 1.67 |
| Validation | | | |
| MolProbity score | 1.11 | 1.71 | 1.85 |
| Clashscore | 1.65 | 4.00 | 4.29 |
| Poor rotamers (%) | 1.20 | 1.85 | 2.82 |
| Ramachandran plot | | | |
| Favored (%) | 97.28 | 95.35 | 95.69 |
| Allowed (%) | 2.72 | 4.65 | 4.31 |
| Disallowed (%) | 0.00 | 0.00 | 0.00 |

| | B2R (Icatibant)-BRIL+SRP2070Fab | | V2R (Tolvaptan)-BRIL |
|---|---|---|---|
| | Overall (EMDB-64536) (PDB 9UVT) | TM focused (EMDB-64537) (PDB 9UVU) | |
| **Data collection and processing** | | | |
| Magnification | 105,000 | | 105,000 |
| Voltage (kV) | 300 | | 300 |
| Electron exposure (e–/Å$^2$) | 45.8, 45.7 | | 47.1 |
| Defocus range (μm) | -0.8 to -1.8, -0.8 to -1.6 | | -0.8 to -1.6 |
| Pixel size (Å) | 0.83 | | 0.83 |
| Symmetry imposed | C1 | C1 | C1 |
| Initial particle images (no.) | 7,108,965 | | 6,751,366 |
| Final particle images (no.) | 187,522 | | 76,583 |
| Map resolution (Å) | 3.29 | 3.69 | 5.95 |
| FSC threshold | 0.143 | 0.143 | 0.143 |
| **Refinement** | | | |
| Initial model used (PDB code) | AF2 model, 7XRZ | AF2 model | |
| Map sharpening *B* factor (Å$^2$) | 126.1 | 141.7 | |
| Model composition | | | |
| Non-hydrogen atoms | 4,529 | 2,209 | |
| Protein residues | 563 | 264 | |
| Ligands | 1 | 1 | |
| *B* factors (Å$^2$) | | | |
| Protein | 137.974 | 117.683 | |
| Ligand | 170.229 | 125.219 | |
| R.m.s. deviations | | | |
| Bond lengths (Å) | 0.009 | 0.009 | |
| Bond angles (°) | 1.65 | 1.70 | |
| Validation | | | |
| MolProbity score | 1.87 | 1.78 | |
| Clashscore | 3.87 | 2.69 | |
| Poor rotamers (%) | 2.82 | 2.87 | |
| Ramachandran plot | | | |
| Favored (%) | 94.90 | 94.49 | |
| Allowed (%) | 5.10 | 5.51 | |
| Disallowed (%) | 0.00 | 0.00 | |

| | V2R (Tolvaptan)-ARK1<br>(EMDB-64541)<br>(PDB 9UVY) | LPA2 (Ki16425)-ARK1<br>(EMDB-64542)<br>(PDB 9UVZ) |
|---|---|---|
| **Data collection and processing** | | |
| Magnification | 105,000 | 105,000 |
| Voltage (kV) | 300 | 300 |
| Electron exposure (e–/Å$^2$) | 45.4 | 45.9, 57.6 |
| Defocus range (μm) | -1.0 to -2.0 | -0.8 to -1.8, |
| Pixel size (Å) | 0.83 | 0.83 |
| Symmetry imposed | C1 | C1 |
| Initial particle images (no.) | 6,079,711 | 12,736,611 |
| Final particle images (no.) | 201,250 | 151,855 |
| Map resolution (Å) | 2.98 | 2.99 |
| FSC threshold | 0.143 | 0.143 |
| **Refinement** | | |
| Initial model used (PDB code) | AF2 model | AF2 model |
| Map sharpening *B* factor (Å$^2$) | 82.2 | 79.2 |
| Model composition | | |
| Non-hydrogen atoms | 5,247 | 5,487 |
| Protein residues | 653 | 682 |
| Ligands | 1 | 1 |
| *B* factors (Å$^2$) | | |
| Protein | 110.738 | 79.240 |
| Ligand | 65.353 | 94.110 |
| R.m.s. deviations | | |
| Bond lengths (Å) | 0.006 | 0.006 |
| Bond angles (°) | 1.67 | 1.80 |
| Validation | | |
| MolProbity score | 1.40 | 1.47 |
| Clashscore | 1.87 | 2.41 |
| Poor rotamers (%) | 1.42 | 2.52 |
| Ramachandran plot | | |
| Favored (%) | 95.21 | 97.21 |
| Allowed (%) | 4.79 | 2.79 |
| Disallowed (%) | 0.00 | 0.00 |
